# Sensory expectations and prediction error during feedback control in the human brain

**DOI:** 10.64898/2026.01.19.700321

**Authors:** Marco Emanuele, Paul L. Gribble, J. Andrew Pruszynski, Jonathan A. Michaels, Jörn Diedrichsen

## Abstract

External disturbances to the body are better counteracted when their nature can be predicted in advance. Here, we investigated the neural mechanisms through which probabilistic predictions shape feedback responses using functional magnetic resonance imaging (fMRI) in humans. We show that, prior to a mechanical perturbation applied to a finger, the primary motor (M1) and somatosensory (S1) cortices receive a signal that linearly encodes the expected sensory input. When perturbations reach these areas, expectations are combined with the sensory input through a simple additive mechanism, yielding motor commands that reflect a weighted sum of the two signals. At the same time, M1 and S1 receive a prediction error signal encoding the difference between expectations and actual sensory input. This signal is visible in fMRI data in humans and in the local field potentials in non-human primates, but not in M1 or S1 spiking activity.

## Main

From skiing to cycling on uneven ground, many activities require rapid compensations for external disturbances to the body. Yet, feedback responses to these mechanical perturbations would arrive too late if exclusively driven by delayed^1–3^ sensory input. Performance can be improved if the nature (e.g., direction, intensity) of upcoming perturbations is predicted from contextual cues or prior experience. For example, a passenger standing on a bus can predict being pulled to the right when the bus is about to turn left at a light. However, if the driver suddenly swerves right to avoid a pedestrian, the passenger will be pushed left, having to reverse the expected response. Preparing movements in advance typically improves performance^4–9^, but it is unclear how the brain shapes feedback responses based on probabilistic knowledge of future perturbations.

Recent work from our group showed that both humans and non-human primates were able to counter elbow perturbations (flexion or extension) more quickly when receiving valid probabilistic information about the upcoming perturbation direction^9^. In monkeys, the expected direction was probabilistically represented in the spiking activity of neurons in the dorsal premotor (PMd) and primary motor cortex (M1). At perturbation onset, these sensory expectations shaped the initial response, which was then increasingly dictated by the accumulating sensory input that signalled the actual perturbation direction. These findings point to a simple mechanism whereby cortical motor regions are pre-activated based on the expectation and then generate the feedback response by additively combining that expectation with the incoming sensory information.

Here, we used 7T functional magnetic resonance imaging (fMRI) to understand how such sensory expectations are encoded in the human brain. While lacking single-neuron and millisecond resolution, fMRI offers broader spatial coverage of cortical motor regions. In addition, the blood-oxygen-level-dependent (BOLD) signal mostly reflects the synaptic input to neural populations^10–12^ and therefore provides a complementary measure to spiking activity. These properties enabled us not only to generalise our results from the monkey to the human brain, but also to gain a deeper understanding of how motor circuits perform the computations necessary for using probabilistic predictions in feedback control. In particular, predictive coding theories posit that the brain computes a prediction error signal, i.e., the difference between the expected and incoming sensory input^13–16^. An explicit representation of this information, however, was surprisingly absent in the spiking activity recorded in monkeys^9^. We reasoned that fMRI may unveil this previously undetected prediction error signal either because it occurs in brain regions not covered by our electrophysiological recordings, or because it is transmitted in the synaptic input to cortical motor regions without directly influencing the spiking activity. To test the latter hypothesis, we systematically compared our fMRI results in humans to both the spiking activity^9^ and the simultaneously recorded local field potentials (LFPs) from analogous areas in monkeys. The LFPs mostly reflect synchronised synaptic input to cortical areas^17^ and are therefore a better predictor of the BOLD signal than spiking activity.

## Results

### Sensory expectations bias responses to mechanical perturbations in a finger perturbation task

To adapt our previous probabilistic perturbation paradigm^9^ for fMRI, we chose a task in which human participants (Experiment 1, N=14; Experiment 2, N=10) countered sudden mechanical perturbations (∼3.5N) to their right index or ring finger (Fig. 1a). Compared to elbow perturbations, finger perturbations are easier to deliver without causing motion artifacts in fMRI data. More importantly, at the relatively low spatial resolution of fMRI, finger representations are more spatially distinct than movement directions^18^, enabling a clearer differentiation of the BOLD activity patterns associated with the two perturbations.

**Figure 1.**
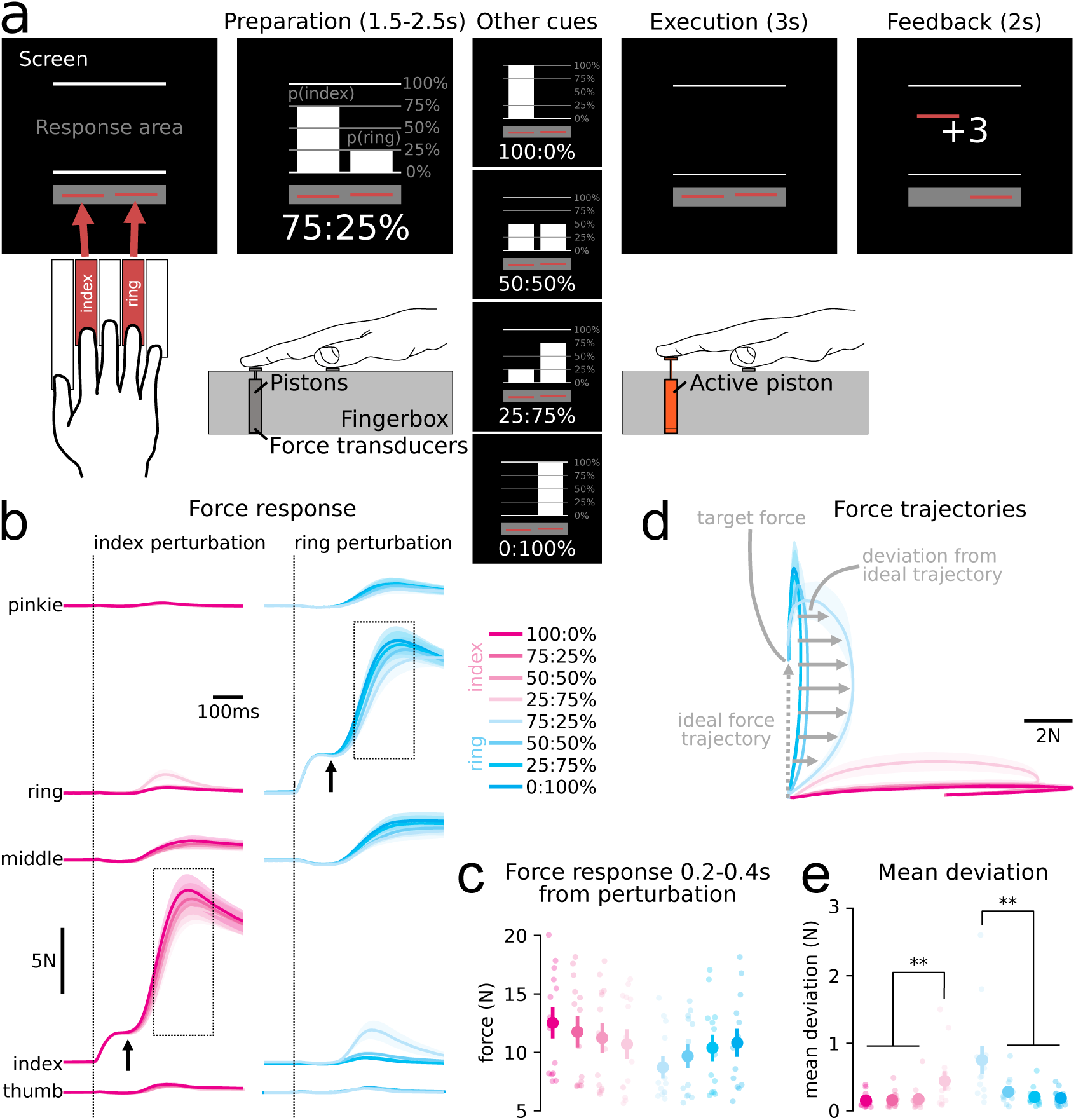
**(a)** Participants placed their right hand on a 5-finger keyboard. During preparation, a visual cue indicated the probability that either the index or ring finger would be perturbed. Participants were instructed to maintain the force on both fingers (indicated by red line cursors) within a 0.1-0.6N range as indicated by the hold area (grey rectangle). During execution, participants had to counter the perturbation applied to the index or ring finger by pushing back the pneumatic piston as quickly as possible. In the feedback phase, participants received a score (−1, 0, +1, or +3) depending on their performance (see Methods). **(b)** Force response of all 5 fingers for index and ring finger perturbations, depending on the cued probability. The vertical dotted line indicates the onset of the mechanical perturbation applied to the index (left column) or ring (right column) finger. The black arrows mark the start of the active force responses. **(c)** Mean force responses between 0.2-0.4s (see dotted rectangles in b) from the perturbation. Dots show individual participants. Error bars denote ±SEM across participants. **(d)** Mean force trajectories for index and ring finger perturbation between 0-0.5s from the perturbation. To assess corrections in finger selection, we calculated, in each trial, the mean deviation from the ideal straight force trajectory (dashed grey arrow). **(e)** The mean deviation was significantly larger when the stimulated finger was the one cued with lower probability. Asterisks denote statistical significance (*P<0.05, **P<0.01, ***P<0.001).

Each trial began with the presentation of a visual cue (preparation phase) signalling the probability that either the index or ring finger would receive the perturbation. After a variable delay of 1.5-2.5s, a perturbation randomly drawn from the cued probability was applied to one of the two fingers using a pneumatic piston. The participant had to respond as quickly as possible by pushing down the piston with the perturbed finger (execution phase).

The active force response began 174±20ms after the perturbation and was modulated by the probability cue (Fig. 1b). Specifically, between 0.2-0.4s after the perturbation, the stimulated finger produced a larger force if cued with higher probability during preparation (Fig. 1c; index, F_3,39_=21.134, P<0.001; ring, F_3,39_=6.109, P=0.002). Often, participants responded first with the finger cued with higher probability even when the perturbation was applied to the other (e.g., 75:25% cue followed by ring perturbation) and then switched to the perturbed finger only after sensory evidence had accumulated. This behaviour is clearly visible in the 2-dimensional force trajectory of the index and ring finger (Fig. 1d). Responding only with the perturbed finger would produce a straight upward or rightward trajectory. Trials in which the perturbation was applied to the finger cued with lower probability showed a significantly larger mean deviation (see Methods) from the ideal straight trajectory compared to those in which the perturbation was applied to the high-probability finger (Fig. 1e; index: t_13_=2.744, P=0.008; ring: t_13_=3.201, P=0.003).

Similar to what we recently reported for the arm^9^, these results show that participants additively combined sensory expectation with the incoming sensory input to generate the initial finger response.

### Representation of expectation and uncertainty during preparation

In Experiment 1, participants performed the task while being scanned with 7T fMRI. To cleanly separate preparation and execution-related activity despite the slow haemodynamic response, we included 2/3 Go- and 1/3 No-Go trials in the design. During No-Go trials, the cue was shown but no perturbation occurred, allowing us to estimate the BOLD response during preparation independent of execution.

BOLD activity within each region-of-interest (ROI; see Fig. 2a) showed a clear separation between Go and No-Go trials after perturbation onset (Fig. 2b). To characterise brain activation, we fit a general linear model (GLM) with separate regressors for preparation and execution (see Methods). During preparation, we observed a significant activation relative to resting baseline in all ROI except S1 (Fig. 2c; Table 1, first row).

**Figure 2.**
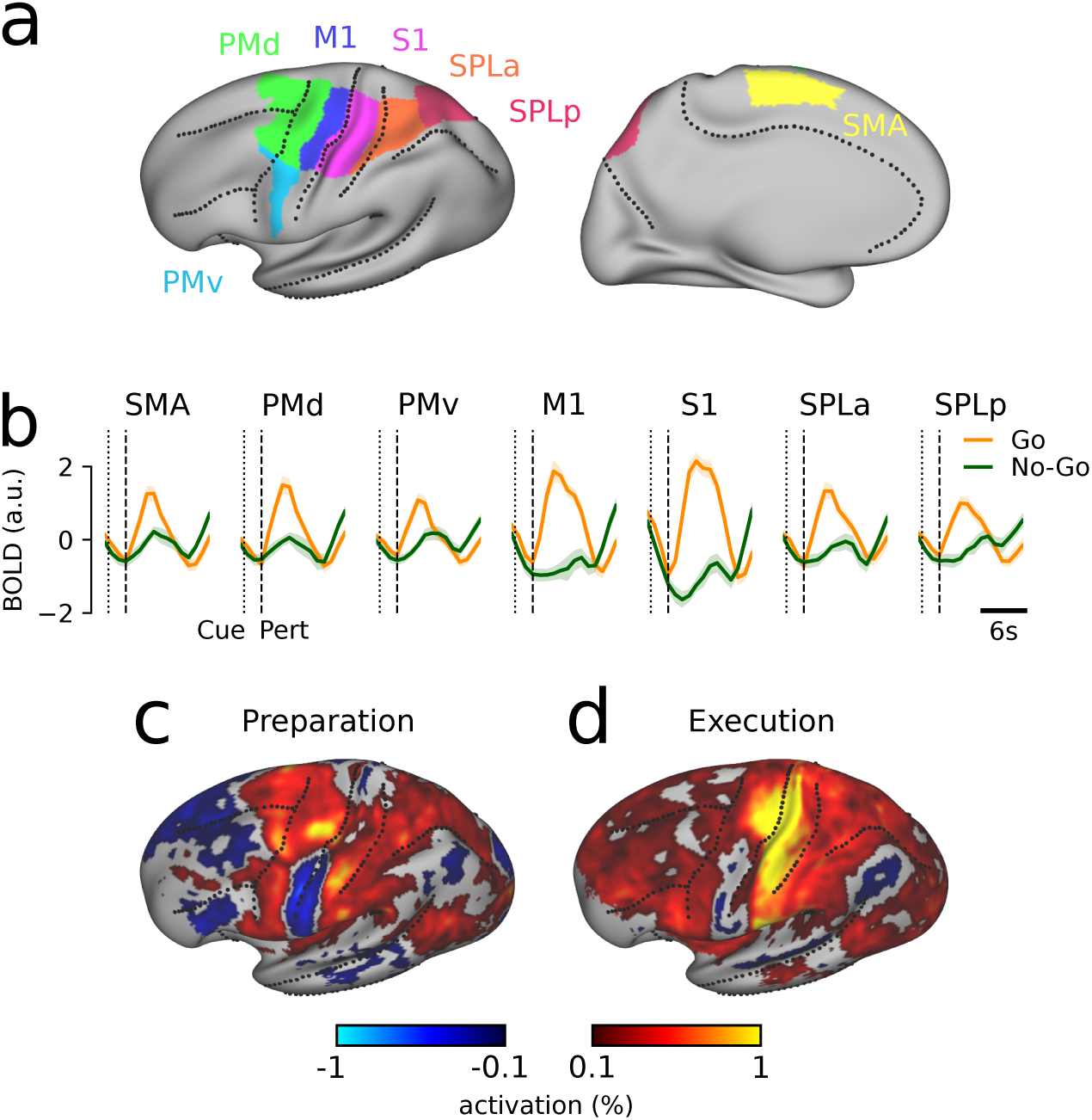
**(a)** ROI as defined on the group inflated surface of the left hemisphere. Dotted lines indicate major sulci. **(b)** BOLD time series averaged across participants in each ROI for Go and No-Go trials, with Go trials time-aligned to perturbation onset (dashed vertical line) and No-Go trials time-aligned to cue onset (dotted vertical line). Error bars indicate ±SEM across participants. **(c-d)** Activation relative to resting baseline (in percent signal change) during (c) preparation and (d) execution in the left hemisphere. Results for the right hemisphere are shown in Supplementary Materials 2.

**Table 1.**
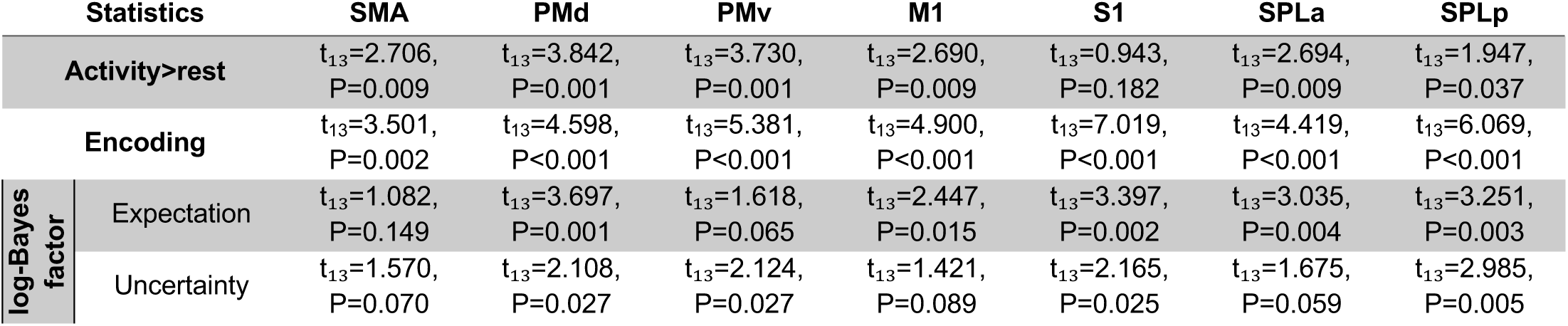
ROI-based statistics for preparation. T-values are one-sided t-tests against 0 with uncorrected P-values provided. For a family-wise error rate of 0.05, the region passes the Bonferroni correction for an uncorrected P-value<0.007.

| Statistics |  | SMA | PMd | PMv | M1 | S1 | SPLa | SPLp |
| --- | --- | --- | --- | --- | --- | --- | --- | --- |
| Activity>rest | | $t_{13}=2.706$,<br>$P=0.009$ | $t_{13}=3.842$,<br>$P=0.001$ | $t_{13}=3.730$,<br>$P=0.001$ | $t_{13}=2.690$,<br>$P=0.009$ | $t_{13}=0.943$,<br>$P=0.182$ | $t_{13}=2.694$,<br>$P=0.009$ | $t_{13}=1.947$,<br>$P=0.037$ |
| Encoding | | $t_{13}=3.501$,<br>$P=0.002$ | $t_{13}=4.598$,<br>$P<0.001$ | $t_{13}=5.381$,<br>$P<0.001$ | $t_{13}=4.900$,<br>$P<0.001$ | $t_{13}=7.019$,<br>$P<0.001$ | $t_{13}=4.419$,<br>$P<0.001$ | $t_{13}=6.069$,<br>$P<0.001$ |
| log-Bayes factor | Expectation | $t_{13}=1.082$,<br>$P=0.149$ | $t_{13}=3.697$,<br>$P=0.001$ | $t_{13}=1.618$,<br>$P=0.065$ | $t_{13}=2.447$,<br>$P=0.015$ | $t_{13}=3.397$,<br>$P=0.002$ | $t_{13}=3.035$,<br>$P=0.004$ | $t_{13}=3.251$,<br>$P=0.003$ |
| | Uncertainty | $t_{13}=1.570$,<br>$P=0.070$ | $t_{13}=2.108$,<br>$P=0.027$ | $t_{13}=2.124$,<br>$P=0.027$ | $t_{13}=1.421$,<br>$P=0.089$ | $t_{13}=2.165$,<br>$P=0.025$ | $t_{13}=1.675$,<br>$P=0.059$ | $t_{13}=2.985$,<br>$P=0.005$ |

Yet, univariate activation offers only a superficial view of the neural processes that occur in a brain region. Even when different conditions drive a region into distinct neural states, activations and deactivations across voxels can cancel out, yielding similar regional averages. We therefore used representational similarity analysis (RSA)^19–21^ to characterise the neural representation of the task in each ROI. We first evaluated whether the preparatory multi-voxel activity patterns differed significantly across the 5 probability cues. To this end, we calculated the cross-validated Mahalanobis (crossnobis) dissimilarities^22^ between the activity patterns elicited by different probability cues in each ROI. In a region without cue representation, the average dissimilarity estimate should be zero. In contrast, all ROI showed above-chance encoding (Table 1, second row). To ensure that this cue representation was not caused by contamination with execution-related activity, we replicated this analysis after estimating the preparatory activity from No-Go trials only (Supplementary Materials 3).

We then asked how the probability cues were represented in each ROI. A priori we hypothesised two possible representational geometries (Fig. 3a). The *expectation* representation encodes the upcoming perturbation (index vs. ring). Therefore, the neural activity patterns for the 100:0% should be maximally different from the 0:100% condition, with the other patterns being weighted averages of these extremes. This corresponds to activity patterns that are linearly ordered according to the expected perturbation (Fig. 3b, horizontal axis). In contrast, in the *uncertainty* representation (Fig. 3b, vertical axis) the neural activity patterns for 100:0% and 0:100% are identical (certain perturbation) but maximally different from the 50:50% condition (undetermined perturbation).

**Figure 3.**
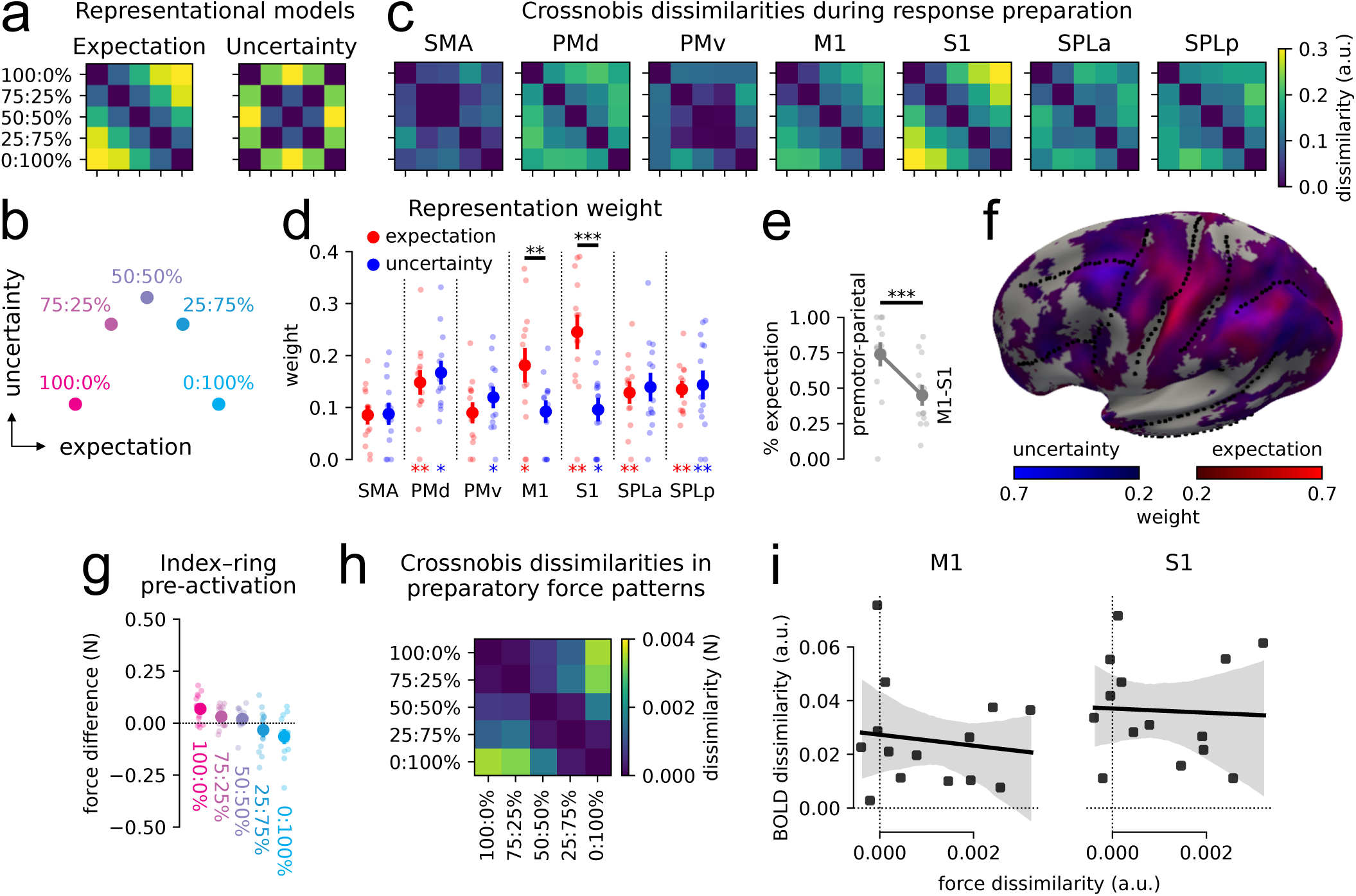
**(a)** Predicted RDMs for the expectation and uncertainty representations. **(b)** Representational geometry of an area with a mixture of expectation and uncertainty encoding. **(c)** Average crossnobis dissimilarities for each ROI. **(d)** Weight of expectation and uncertainty in each ROI. The asterisks below the bars denote the statistical significance (*P<0.05, **P<0.01, ***P<0.001) of the log-Bayes factor against 0, indicating that removing the component significantly reduces the overall model performance (see Methods). Error bars indicate ±SEM across subjects. The horizontal bars denote significant weight differences between expectation and uncertainty. Dots show individual participants. **(e)** Weight of expectation normalized by the sum of expectation and uncertainty in M1-S1 and premotor-parietal regions. **(f)** Weight of expectation and uncertainty component in a continuous searchlight analysis conducted on the surface of the left hemisphere. **(g)** Average difference between the force produced by the index and ring finger during the preparation phase. Dots denote individual participants. Vertical bars indicate ±SEM. **(h)** Crossnobis dissimilarities between force patterns during preparation. **(i)** Linear fit between average crossnobis dissimilarity observed in force and BOLD activity patterns during preparation. The black dots denote individual participants. The shaded region represents the 95% confidence interval (CI) for the regression line.

Visual inspection of the representational dissimilarity matrices (RDMs; Fig. 3c) suggested that the representational geometry in M1 and S1 closely corresponded to a linear encoding of the sensory expectation. In contrast, the RDMs in premotor and parietal areas were more similar to the predicted RDM for the uncertainty representation. We used pattern component modelling (PCM) to quantify these observations and express the information content in each region as a weighted combination of expectation and uncertainty. Both representations contributed to preparatory activity patterns. Indeed, removing either representation worsened the model performance, as indicated by significantly positive log-Bayes factor (see Methods) in most ROI for both expectation and uncertainty (Table 1, third and fourth rows). While this suggests that preparatory activity across cortical motor areas represents both expectation and uncertainty, the strength of the two representations varied markedly.

In M1 and S1, the weight of the expectation representation was significantly higher than that of uncertainty (Fig. 3d; M1, t₁₃=2.934, P=0.006; S1, t₁₃=4.056, P<0.001). By contrast, in premotor and parietal areas this difference was not statistically reliable (all t_13_<1.108, P>0.144). To directly demonstrate the different information in the two groups of regions, we assessed the weight of the expectation relative to the summed weight for both representations (Fig. 3e). The proportional expectation weight was significantly larger in M1 and S1 compared to premotor and parietal areas (t₁₃=4.974, P<0.001). The different weight of expectation and uncertainty in M1-S1 and premotor-parietal areas is also evident in a continuous searchlight analysis (Fig. 3f).

### The expectation representation is not caused by overt motor output during preparation

While the expectation representation in M1 and S1 was strong, we needed to rule out that it was driven by subtle finger pre-activation. To discourage unwanted modulation of finger force before perturbation onset, we required that participants kept the two cursors indicating the force produced by the index and ring finger in a range between 0.1-0.6N (see Fig. 1a). Participants complied well with this instruction, but pressed slightly harder with the pre-cued finger, with an average modulation across index and ring finger between the 100:0% and 0:100% probability cues of 0.133±0.119N (Fig. 3g; F_4,52_=10.220, P<0.001, repeated-measures ANOVA). In line with this effect, crossnobis dissimilarities between the force patterns produced by the 5 fingers during preparation suggested that this subtle finger pre-activation reflected the cued probability (Fig. 3h).

To test whether these pre-activation patterns could explain the expectation representation in M1 or S1, we first performed a linear regression analysis between the average crossnobis dissimilarity observed in the forces (see Fig. 3h) and in the preparatory activity patterns (see Fig. 3c). We found no systematic relationship between these two variables (Fig. 3i; M1, slope=-2.042, P=0.665; S1, slope=-0.786, P=0.869). More importantly, the intercept of the regression (i.e., our estimate of the neural difference for participants that did not show any expectation-driven pre-activation) was significantly larger than 0 both for M1 and S1 (M1, intercept=0.027, P=0.001; S1, intercept=0.037, P<0.001).

In a second control analysis, we regressed out, within each participant, the force dissimilarities (Fig. 3h) for each condition pair from the pairwise BOLD dissimilarities in each ROI (Fig. 3c). The residual BOLD dissimilarities (Fig. S4a) remained significantly positive across all ROI (Table S4, first row), and the comparison with the expectation and uncertainty models revealed similar results to the main analysis (Fig. S4b; see Table S4, second to fifth rows). In conclusion, these two complementary control analyses demonstrate that the expectation representation during preparation is not caused by the pre-activation of the pre-cued finger.

### Electrophysiology: Expectation signal in S1 LFPs but not in spiking activity

In contrast to our current fMRI results, our previous findings in non-human primates did not show a strong expectation signal in S1^9^. To enable a meaningful comparison of our results in humans and monkeys, we revisited our electrophysiological recordings from monkeys performing the arm perturbation task^23^ and used PCM to fit the relative weight of expectation and uncertainty to the spiking activity across cortical motor regions. Unlike in PMd and M1, the expectation representation was nearly absent in S1 (Fig. 4b, red lines; see Supplementary Materials 6 for results about uncertainty).

**Figure 4.**
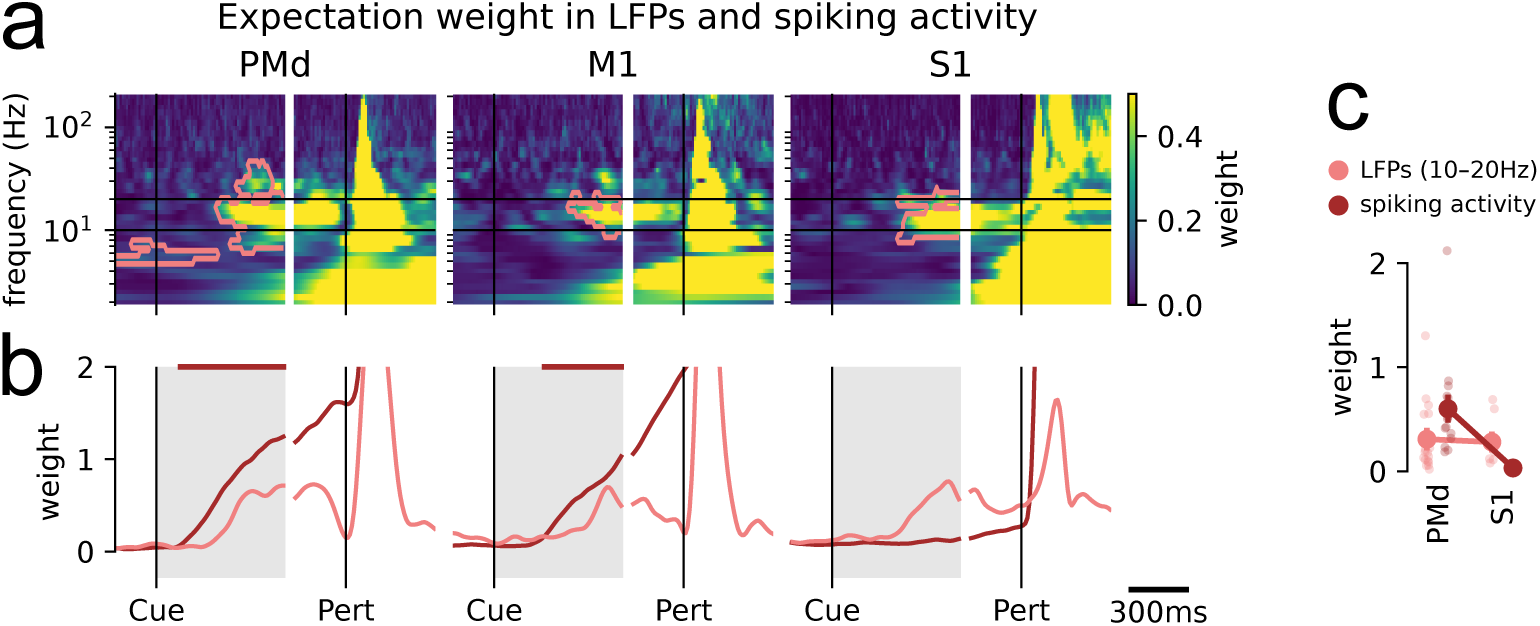
**(a)** Weight of expectation in the LFPs and **(b)** spiking activity (red lines) recorded from PMd, M1 and S1 in our previous non-human primate dataset^9^, aligned to cue presentation (Cue) and perturbation onset (Pert). In panel a, pink contours denote time-frequency clusters where the log-Bayes factor for expectation was significantly higher than 0. In panel b, pink lines show the average weight of expectation in the LFPs within the 10-20Hz frequency interval. **(c)** Average expectation weight in the grey-shaded time interval in panel b. Note that dots refer to individual recording sessions rather than different monkeys.

While this discrepancy could be due to differences between paradigms or species, it could also depend on the physiological underpinnings of the different recording modalities. Spiking activity reflects the neuronal output, whereas the BOLD signal is more influenced by the synaptic input to neural populations^10,11^. To test the idea that expectations are represented in the synaptic input to S1 without influencing the spiking activity, we fit the relative weight of expectation and uncertainty to the LFPs recorded simultaneously with spiking activity in our previous electrophysiological dataset^23^ and reflecting synchronised synaptic input^17^ (see Supplementary Materials 5 for power modulations across frequency bands). We found that, in the LFPs recorded from both PMd and S1, expectations were significantly represented (P<0.05; cluster-based permutations) in a frequency band from 10-20Hz (i.e., low beta-band; Fig. 4a; see Supplementary Materials 6 for results about uncertainty). In these areas, the expectation weight in the LFPs and spiking activity showed a significant interaction between recording modality and ROI (Fig. 4c; F_1,48_=7.761, P=0.008). Thus, a parsimonious explanation for the apparent discrepancy between fMRI and electrophysiological findings is that the information about expectations reaches S1 as synaptic input but does not influence the neuronal spiking.

### Expectations pre-activate the representation of the future sensory input

We then investigated how the expectation patterns were related to the activity patterns elicited by incoming sensory information. To this end, we calculated the difference between the activity patterns in M1 and S1 for index (100:0% and 75:25% cues) and ring finger (0:100% and 25:75% cues) expectation during preparation (i.e., expectation pattern) and the difference between the activity patterns for index and ring perturbation during execution (i.e., sensory input pattern). We then estimated the correlation between the expectation and sensory input patterns. At one extreme, the expectation may simply pre-activate the sensory representation of the most likely finger. In this scenario, the two patterns should correlate perfectly. At the other extreme, the expectation may elicit activity patterns that are orthogonal to those elicited by the sensory input in voxel space. In this case, the expectation and sensory pattern would be uncorrelated.

Hypotheses about the true size of correlations are difficult to test using Pearson’s correlation, because measurement noise always attenuates correlations towards zero. Consequently, even if the true underlying pattern is the same, Pearson’s correlation will be lower than 1. We therefore estimated correlations using a maximum-likelihood approach that corrects for this attenuation bias by estimating measurement noise from repeated samples of the expectation and sensory input patterns in different runs (see Methods)^24^.

In M1 and S1, the estimates of the true correlation between preparation and execution patterns were positive but smaller than 1 (Fig. 5; M1, ρ=0.632, 95% CI [0.505, 0.711]; S1, ρ=0.611, 95% CI [0.487, 0.718]). Therefore, the expectation of a perturbation on a specific finger pre-activates the voxels that also receive the incoming sensory information caused by the perturbation. At the same time, a correlation of ∼0.6 implies that roughly two thirds of the variance of the execution pattern is not anticipated by preparatory activity, suggesting that in these two phases cortical activity explores overlapping, but slightly separate subspaces.

**Figure 5.**
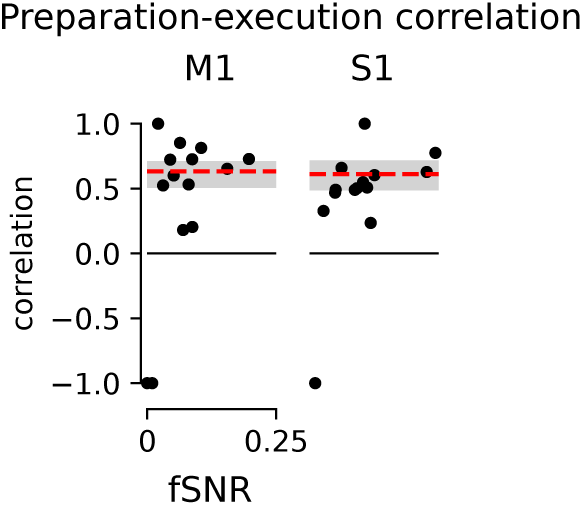
Maximum-likelihood correlation estimates between preparatory and execution activity patterns. The correlation estimates for each participant are plotted against the ratio of signal variance and noise variance (functional signal-to-noise ratio, fSNR; see Methods). The dashed red line corresponds to the group correlation estimate. The shaded grey area denotes the 95% central CI established by a participant-wise bootstrap procedure. Note that, when the fSNR approaches zero, the correlation estimate is unreliable and the maximum likelihood converges towards +1 or −1, depending on the sign of the covariance between the two variables.

### Representation of sensory input and surprise during execution

We then asked how the expectation signal is combined with the incoming sensory information during the execution phase. The mechanical perturbation applied to the fingertip and the ensuing participant’s response elicited a strong increase in the BOLD signal across cortical motor areas (see Fig. 2b,d). To examine the representational geometry during execution, we computed the crossnobis dissimilarities between activity patterns elicited by index or ring finger stimulation. For each finger, we also split the data by the cued probability. We hypothesised that representational geometries would have three components (Fig. 6a). First, execution activity should reflect the *sensory input* (i.e., the stimulated finger). Second, the *expectation* representation identified in preparatory activity may be sustained into the execution phase independently (see Methods) of the sensory input representation. Finally, the brain may represent how *surprising* the perturbation was relative to the expectation, i.e., a representation of the absolute prediction error between expectation and sensory input. Using PCM, we estimated the relative weights for the sensory input, expectation and surprise representations.

**Figure 6.**
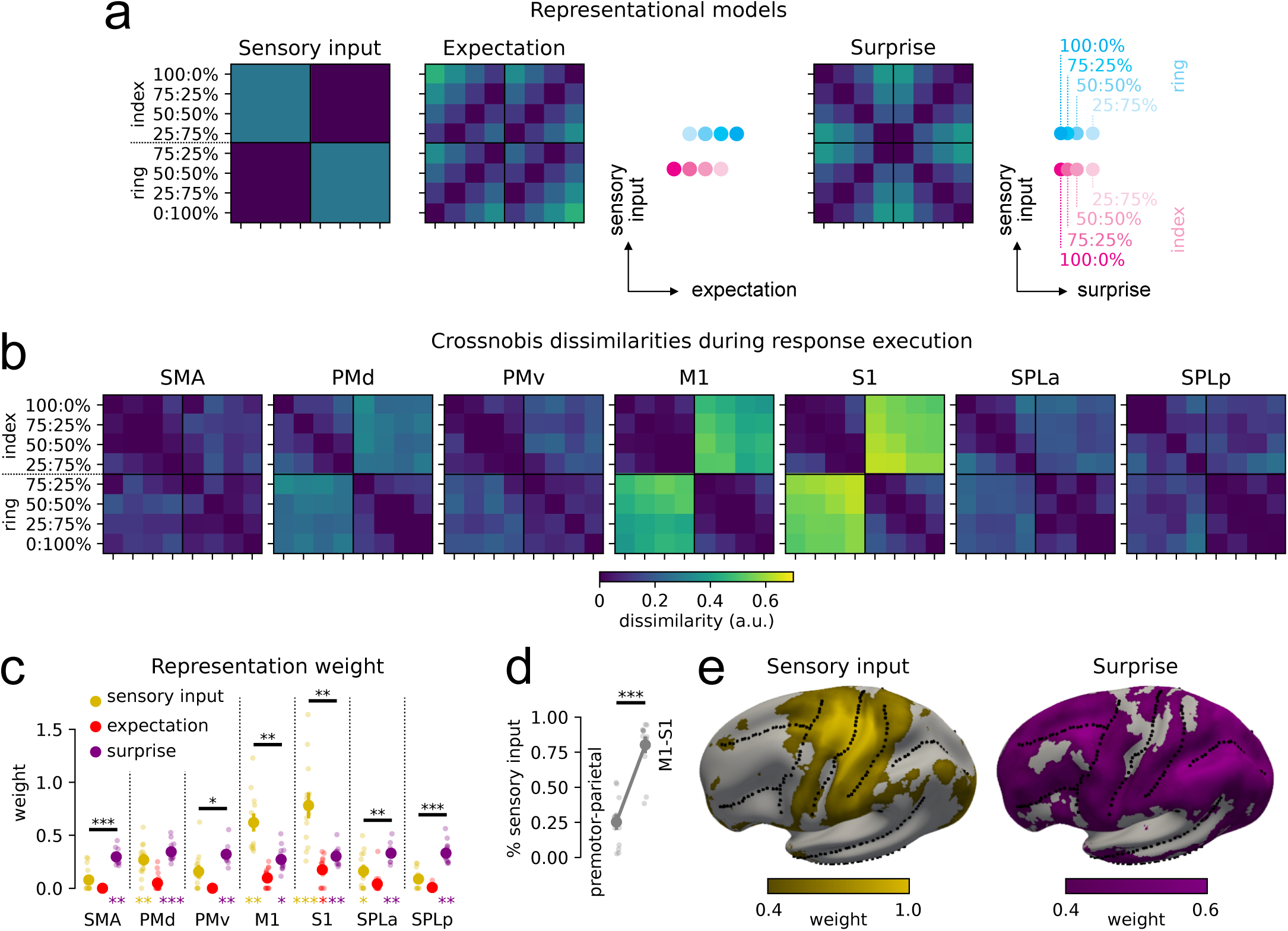
**(a)** Predicted RDMs for sensory input, expectation and surprise and corresponding representational geometries during the execution phase. **(b)** Crossnobis dissimilarity during execution. Note that the representational geometry in M1 and S1, besides a strong effect of the stimulated finger, also reflects the negative correlation model in Fig. 7d. For comparison, see EMG activity in the LLR and Vol time windows in Fig. 7e, which instead reflects the positive correlation model in Fig. 7b. **(c)** Representation weight of sensory input, expectation and surprise in each ROI. The asterisks below the bars denote that the log-Bayes factor of the corresponding representation was significantly larger than 0 (*P<0.05, **P<0.01, ***P<0.001). The horizontal bars with asterisks denote a significant difference between sensory input and surprise. Dots show individual participants. **(d)** Relative weight of sensory input vs. sensory input+surprise in primary sensorimotor and premotor-parietal regions. The horizontal bar with asterisks denotes the significant difference between the two groups of ROI. **(e)** Representation weight of sensory input and surprise projected on the inflated surface of the left hemisphere.

The empirical RDMs suggested a strong representation of the sensory input, especially in M1 and S1 (Fig. 6b). The log-Bayes factor (see Methods) for sensory input was significantly positive in PMd, M1, S1 and SPLa (Table 2, third row). Furthermore, all ROI showed a significant representation of surprise (Table 2, fifth row). In contrast, no expectation representation independent of the sensory input was found during the execution phase in any ROI except S1 (Table 2, fourth row).

**Table 2.** ROI-based statistics for execution. See Table 1 for statistical tests and conventions.

| Statistics |  | SMA | PMd | PMv | M1 | S1 | SPLa | SPLp |
| --- | --- | --- | --- | --- | --- | --- | --- | --- |
| Activity>rest | | $t_{13}=2.614$,<br>P=0.011 | $t_{13}=4.025$,<br>P=0.001 | $t_{13}=2.686$,<br>P=0.009 | $t_{13}=6.912$,<br>P<0.001 | $t_{13}=16.411$,<br>P<0.001 | $t_{13}=6.003$,<br>P<0.001 | $t_{13}=6.163$,<br>P<0.001 |
| Encoding | | $t_{13}=3.741$,<br>P=0.002 | $t_{13}=4.633$,<br>P<0.001 | $t_{13}=4.872$,<br>P<0.001 | $t_{13}=4.856$,<br>P<0.001 | $t_{13}=4.096$,<br>P=0.001 | $t_{13}=4.644$,<br>P<0.001 | $t_{13}=5.413$,<br>P<0.001 |
| log-Bayes factor | Sensory input | $t_{13}=-0.437$,<br>P=0.665 | $t_{13}=2.812$,<br>P=0.007 | $t_{13}=1.346$,<br>P=0.101 | $t_{13}=3.751$,<br>P=0.001 | $t_{13}=5.743$,<br>P<0.001 | $t_{13}=2.111$,<br>P=0.027 | $t_{13}=-0.751$,<br>P=0.767 |
| | Expectation | $t_{13}\ll 0$,<br>P=1.000 | $t_{13}=-4.487$,<br>P=1.000 | $t_{13}\ll 0$,<br>P=1.000 | $t_{13}=-0.860$,<br>P=0.797 | $t_{13}=2.429$,<br>P=0.015 | $t_{13}=-1.262$,<br>P=0.885 | $t_{13}\ll 0$,<br>P=1.000 |
| | Surprise | $t_{13}=3.482$,<br>P=0.002 | $t_{13}=4.817$,<br>P<0.001 | $t_{13}=3.224$,<br>P=0.003 | $t_{13}=2.415$,<br>P=0.016 | $t_{13}=2.906$,<br>P=0.006 | $t_{13}=3.695$,<br>P=0.001 | $t_{13}=2.692$,<br>P=0.009 |

The relative strength of the sensory input and surprise representations, however, differed across ROI. The sensory input was prominent in M1 and S1, with a significantly larger weight than surprise (Fig. 6c; M1, t₁₃=3.422, P=0.002; S1, t₁₃=3.347, P=0.003). In contrast, the surprise representation was significantly stronger than the sensory input in most premotor and parietal areas (SMA, t₁₃=4.816, P<0.001; PMv, t₁₃=1.819, P=0.046; SPLa, t₁₃=3.034, P=0.005; SPLp, t₁₃=4.872, P<0.001). The proportion of sensory input weight relative to the summed weight of sensory input and surprise was significantly larger in M1-S1 compared to premotor-parietal areas (Fig. 6d; t₁₃=14.686, P<0.001), providing direct support to the distinct representation of these two features across different ROI. The different distribution of sensory input and surprise representations can also be seen in a continuous searchlight analysis (Fig. 6e).

### Integration of expectation and sensory input across the motor hierarchy

While the expectation was not represented during execution independently of the perturbation, the modulation of the force response depending on the cued probability (see Fig. 1b,c) suggests that participants integrated the expectation with the incoming sensory information. What form does this integrated signal take in execution activity?

One possibility is that expectation and sensory input are combined additively, with execution activity corresponding to the weighted sum of the two signals. In a region that performs this operation, the expectation and sensory input would drive execution activity patterns in the same direction (Fig. 7a). This would predict that activity patterns for index and ring finger stimulation are more dissimilar to each other if the perturbation is preceded by a congruent as compared to an incongruent cue (Fig. 7b). In this case, the pattern components for expectation and sensory input in execution activity should be positively correlated. Alternatively, the brain could calculate a weighted difference between expectation and sensory input, yielding a signed prediction error signal. In this case, the expectation and sensory input would drive activity patterns in opposite directions (Fig. 7c). This predicts that activity patterns for index and ring stimulation should be more dissimilar if the perturbation is preceded by an incongruent cue (Fig. 7d). Consequently, the correlation between the two pattern components should be negative.

**Figure 7.**
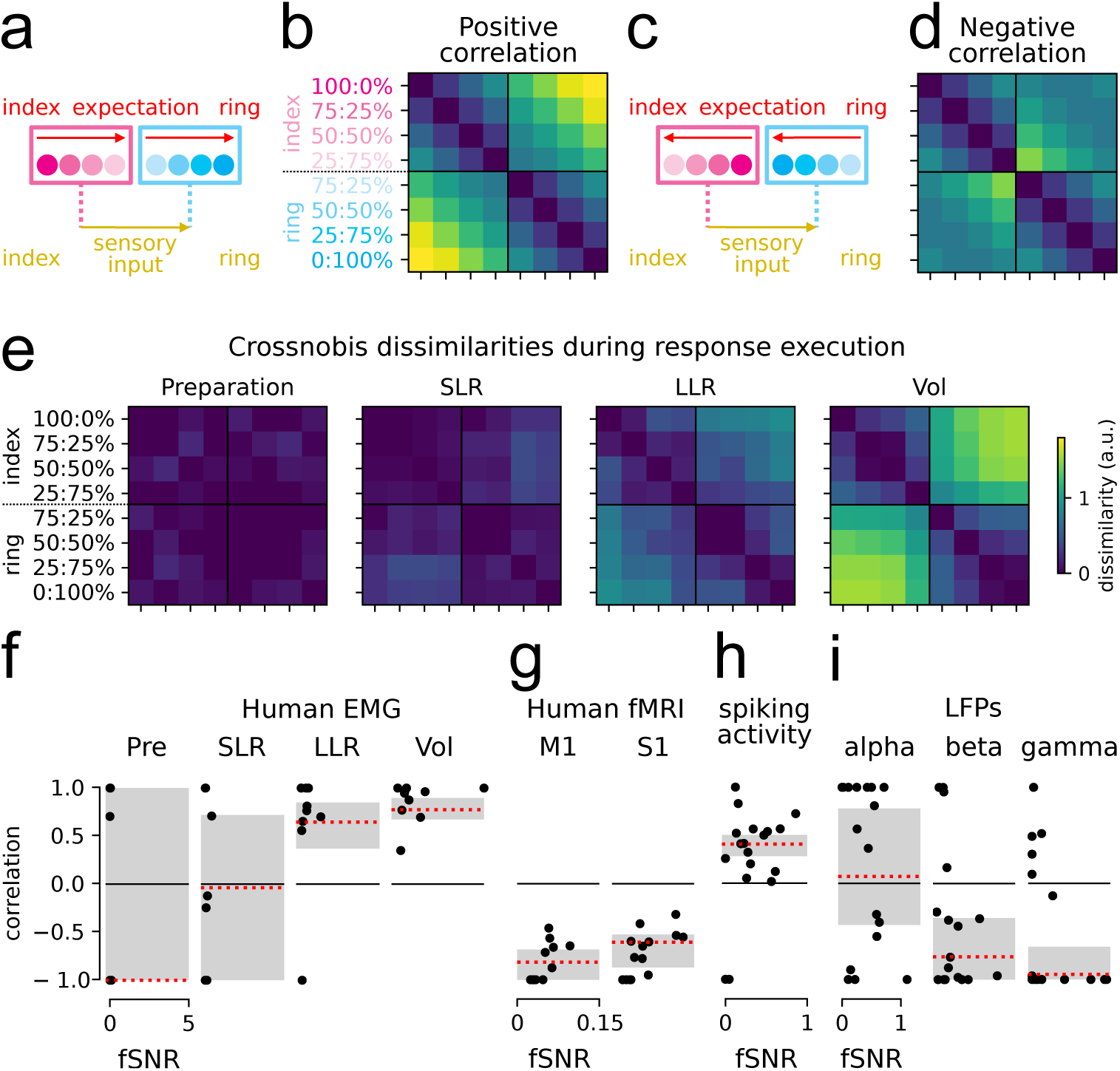
**(a-d)** Representational geometries and predicted RDMs for (a,b) positive and (c,d) negative correlation between expectation and sensory input. The arrows in panels a and c indicate the direction of ring finger encoding in the expectation and sensory input pattern components during execution. In panel a (positive correlation), ring expectation (red arrow) drives activity patterns in the same direction as ring perturbation (yellow arrow). In panel c (negative correlation), ring expectation drives activity patterns in the opposite direction with respect to ring perturbation. **(e)** Average crossnobis dissimilarities between EMG activity patterns across participants. **(f-i)** Maximum-likelihood correlation estimates between expectation and sensory input in (f) EMG activity, (g) estimated BOLD activity patterns from human participants, and (h) spiking activity and (i) LFPs in M1-S1 in non-human primates. The black dots indicate individual participants in panels f and g, and different recording sessions in panels h and i.

We first examined how expectation and sensory input are combined in the motor output. The stronger force responses for expected than unexpected perturbations (see Fig. 1b,c) already suggest that in the motor output expectations and sensory input are combined additively. To characterise the motor output in more detail, in Experiment 2 we recorded the electromyographic (EMG) activity of hand muscles (see Methods) while participants performed the task seated at a desk outside of the scanner. We then used PCM to estimate the correlation between the pattern components for expectation and sensory input elicited in EMG activity after the perturbation. There was no clear modulation in the 100-ms time window (Pre) before the perturbation (Fig. 7e,f; no reliable fSNR, see Methods), nor in the short-latency reflex (SLR; 25-50ms from perturbation; ρ=-0.036, 95% CI [−1.000, 0.719]). This is expected because SLRs are entirely mediated by a spinal circuit^25^ and typically show limited modulation based on contextual influences^26,27^. On the other hand, we found a positive correlation between expectation and sensory input in the long-latency reflex (LLR; 50-100ms; ρ=0.646, 95% CI [0.370, 0.850]) and in the voluntary response (Vol; 100-500ms; ρ=0.774, 95% CI [0.673, 0.896]), consistent with the notion that fast transcortical feedback loops are subject to more sophisticated modulations compared to spinal circuits, and similar to slower volitional responses^28^.

In contrast to EMG activity (Fig. 7e), the RDMs estimated from BOLD activity patterns in M1 and S1 during the execution phase (Fig. 6b) were more similar to the predicted RDM under the hypothesis that the area computes a prediction error signal (Fig. 7c). Confirming this observation, we found a negative correlation between the BOLD patterns for expectation and sensory input in M1 and S1 during execution (Fig. 7g; M1, ρ=-0.817, 95% CI [−1.000, −0.685]; S1, ρ=-0.609, 95% CI [−0.872, −0.529]). Therefore, the BOLD activity in M1 and S1 during execution did not reflect the weighted additive integration of the expected and actual sensory input, as observed in the EMG, but the difference between the two signals, i.e., a signed prediction error signal.

### Electrophysiology: Prediction error in M1-S1 LFPs but not in spiking activity

In contrast to our fMRI results, our previous analysis of the electrophysiological dataset in non-human primates did not show a signed prediction error signal in either M1 or S1^9^. We therefore reanalysed these data using the same methods as for our fMRI data, and again both in the spiking activity and in the LFPs. The spiking activity of neurons in M1 and S1 showed a positive correlation between expectation and perturbation direction (Fig. 7h; ρ=0.407, 95% CI [0.280, 0.503]) from 0.04-0.24s after the perturbation, similar to EMG activity in the human dataset, and consistent with an additive integration. However, as observed in the BOLD signal in human participants, in the same time window the LFPs showed a negative correlation (Fig. 7i) both in the beta (13-25Hz; ρ=-0.763, 95% CI [−1.000, −0.359]) and gamma (25-100Hz; ρ=-0.947, 95% CI [−1.000, −0.659]) frequency bands.

Together, these findings indicate that the synaptic input in M1 and S1, both in monkeys and humans, reflects the signed prediction error between the expected and the actual perturbation. In contrast, the spiking activity in both areas corresponds to a signal that additively combines expectations and perturbation and can be used to drive the muscles.

### Summary of results

To summarise our fMRI results in humans, we projected the BOLD activity patterns for each condition onto the dimensions in voxel space that explained most variance across conditions (Fig. 8a). The first principal component (PC1) reflected the difference between preparation and execution. The second principal component (PC2) reflected the dimension that represents the expected (during preparation) or perturbed (during execution) finger. Along this dimension, the preparatory activity patterns from M1 and S1 (Fig. 8a, magentacyan gradient dots) were linearly ordered according to the expected finger (see Fig. 3c). This ordering was consistent with the direction in which sensory input drove the activity patterns during the execution phase, in line with a positive correlation between preparatory and execution activity (see Fig. 5). For example, expecting a perturbation on the index finger (magenta, 100:0%) pushed preparatory activity patterns to the left along PC2, the same direction as the later index finger perturbation. Instead, expecting (cyan, 0:100%) and responding to a ring finger perturbation pushed the neural activity patterns in the opposite direction.

**Figure 8.**
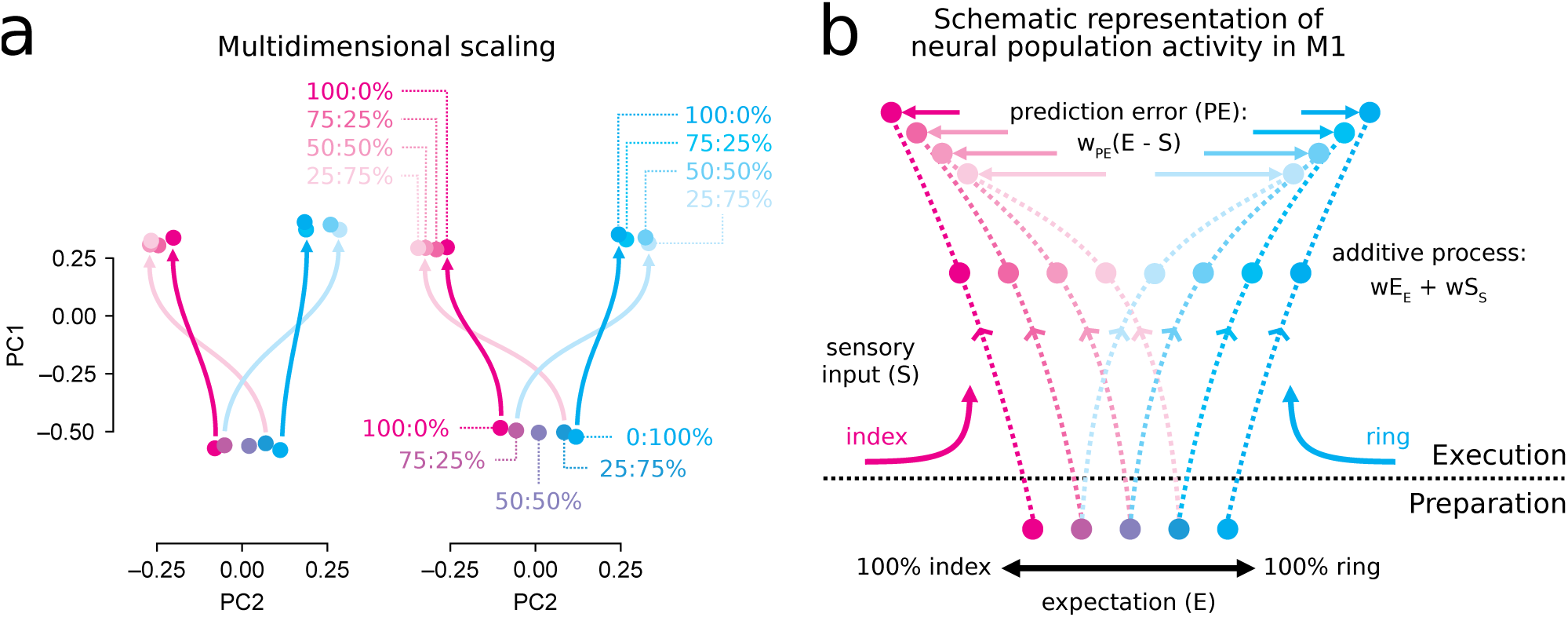
**(a)** Projections of M1 and S1 BOLD activity patterns during preparation and execution onto the two dimensions in voxel space that explained most variance across conditions. The dots along the magenta-cyan gradient correspond to the five probability cues. The magenta- and cyan-shaded dots reflect the activity patterns for index and ring perturbation, respectively. The darker magenta and cyan arrows show the transition from preparatory activity for the 100:0% and 0:100% cues to execution activity for the cued finger. The pink and light blue arrows show the transition from preparatory activity for the 25:75% and 75:25% cues to execution activity for the unexpected finger. The cross-over of the magenta-pink and cyan-light blue arrows highlights the encoding of the prediction error during execution. **(b)** Schematic representation of the integration of expectation and sensory input in M1 spiking activity. During preparation, neural population activity reflects the expectation about the upcoming perturbations (see Fig. 4b). During execution, the expectation is combined with the incoming sensory input through a simple additive mechanism (see Fig. 7h). Additionally, M1 receives a prediction error signal (horizontal magenta and cyan arrows) that may push the execution activity towards a state corresponding to the actual perturbation or could be used for the updating of expectations during learning.

After the perturbation, the BOLD activity patterns were more dissimilar when the stimulated finger was cued with lower probability (Fig. 8a, pink and light blue dots). This is consistent with expectation and sensory input pushing execution activity patterns in opposite directions, resulting in a negative correlation between the two pattern components that is indicative of the computation of a prediction error signal (see Fig. 7g). For example, a highly expected perturbation to the index finger (e.g., following a 100:0% cue) pushed execution activity to the right along PC2, that is, toward activity patterns elicited by a ring perturbation. On the other hand, an unexpected perturbation to the index finger (e.g., following a 25:75% cue) pushed execution activity to the left along PC2, away from activity patterns elicited by a ring perturbation.

## Discussion

Our fMRI results in humans demonstrate that BOLD activity patterns in PMd and M1 scale linearly with sensory expectations about upcoming finger perturbations. These results are consistent with our recent findings in non-human primates showing that spiking activity in PMd and M1 linearly represents expectations about upcoming elbow perturbations (see Fig. 8b, preparation)^9^. In the fMRI data, we found a similar expectation representation in S1, which was absent in the spiking activity recorded in the monkeys. This discrepancy likely occurs because the BOLD signal is more related to the synaptic input to neural populations and less to spiking activity itself^10,11^. Indeed, the LFPs recorded from non-human primates also showed a strong expectation signal in S1. Together, these results suggest that both M1 and S1 receive information about expectations in their synaptic input. Yet, unlike in M1, in S1 this input information does not influence the local spiking activity. This is consistent with the notion that the spiking of S1 neurons is tightly linked to the actual sensory input^29^, without an expansive representation of other latent dimensions as in M1^30–32^. Interestingly, previous studies have shown similar expectation signals in the BOLD signal recorded from S1 during the preparation of self-initiated movements^33–36^. This suggests that the expectation signal elicited by probabilistic information taps into a more general mechanism that injects information about an upcoming movement in the synaptic input to S1 before its onset.

Whether the expectation signal in S1 serves a specific function remains an open question. One possibility is that this signal modulates or improves subsequent sensory processing in S1^37^. On the other hand, our previous work shows that S1 spiking activity is not modulated by expectations during execution (different from M1)^9^. Therefore, it remains possible that the expectation signal in the synaptic input to S1 is epiphenomenal, and reflects the transmission of this information from other motor regions (e.g., M1, PMd) through corticocortical projections^38–40^.

In both M1 and S1, the BOLD activity patterns during preparation and execution were positively correlated, which indicates that the neural populations pre-activated during preparation are those that receive sensory information from the finger cued with higher probability. Once the perturbation began, the expectation was combined additively with the incoming sensory input. In the finger task, this was visible from the stronger force response when the perturbation was applied to the finger cued with higher probability. The same finger dominated the initial force response even when the perturbation violated the expectations. This is consistent with our previous findings in monkeys showing that the expected feedback response is triggered by an unspecific signal that marks the beginning of the perturbation, but contains little information about its direction^9^. The expectation signal would bias the initial feedback response triggered by this unspecific signal towards the expected finger. In line with this idea, the spiking activity of M1 neurons recorded in monkeys, as well as the initial EMG response in human participants, reflected a weighted sum of expectation and sensory input (see Fig. 8b, execution).

In contrast, the synaptic input in M1 and S1, as indexed by BOLD or LFPs, reflected the signed prediction error (i.e., the difference) between expectation and sensory input. This is surprising because, from a control perspective, the nervous system would not need to compute the prediction error. The weighted sum of expectation and sensory input would be sufficient to guide feedback responses and can be computed without the explicit representation of their difference.

There are two potential, not mutually exclusive, uses for this signed prediction error signal. First, the mismatch between expectation and sensory input could be added to the response, driving the system toward the correct response more rapidly than the sensory input alone (Fig. 8b, horizontal magenta and cyan arrows). This is consistent with predictive coding theories of motor control, which propose that motor commands are adjusted based on the difference between expectations and incoming sensory information^13,14,16,41^. Second, the signed prediction error could be used for updating the expectation for subsequent trials^42^. This updating signal could be weighted by the precision of the expectation^15^, reflected in the uncertainty representation we observed in premotor and parietal regions.

Where could the signed prediction error be computed? The absence of prediction error signals in the spiking activity of M1 and S1 makes it unlikely that this subtractive operation is performed locally. More likely candidates are premotor and parietal regions. These regions receive information about both expectations and sensory input, although not as strongly as M1 and S1. The fMRI data show that this information is combined to calculate the unsigned prediction error, i.e., the surprise representation. Assuming independent and spatially intermingled neural populations encoding the signed prediction error separately for each finger, pooling their activity in the BOLD signal would rectify the signal and produce a response that reflects only the magnitude of the mismatch. At the same time, the original, unrectified prediction error signal transmitted to M1 and S1 could contribute to the representational geometry we found in the synaptic input to these regions. Alternatively, it is also possible that the prediction error signal is computed in subcortical structures, such as the cerebellum^43,44^.

In conclusion, we show that the synaptic input to M1 and S1 is similarly modulated in humans and monkeys responding to sudden mechanical perturbations applied to the fingers or the arm, respectively. These results provide new insights into the neural machinery that governs rapid feedback responses. Expectations are not only represented in the expansive latent dimensions of M1^30,31^ without causing overt muscle activity^9^, but are also evident in S1’s synaptic input. As perturbations occur, feedback responses may not only benefit from the additive combination of this expectation signal with the incoming sensory information but also be further refined by the prediction error signal acting as a corrective drive in the synaptic input to M1.

## Methods

### Participants

We recruited 14 participants for Experiment 1 (age 18-34 years, mean 21.35 years, SD 3.77 years; 6 females) and 10 participants for Experiment 2 (age 21-32 years, mean 25.70 years, SD 4.16 years; 2 females). All participants were right-handed and did not report any neurological condition. The experimental procedures were approved by the Research Ethics Committee at Western University (HSREB 107061 for Experiment 1 and HSREB 108479 for Experiment 2). Participants provided written informed consent and were compensated for their participation.

### Apparatus

We used a custom-made MRI-compatible keyboard device to deliver mechanical perturbations independently to the right index or ring fingertip and record the force response generated by the stimulated finger (Fig. 1a). The keys were equipped with force transducers that measured the isometric force generated by each finger (Honeywell FS series; dynamic range 0–16N; sampling frequency 500Hz). The fingers were comfortably restrained by a padded clamp adjusted to each participant’s hand size. The mechanical perturbation (∼3.5N) was delivered using pneumatic pistons (diameter 3mm) embedded underneath each key and operated by compressed air (∼70psi).

### Task

Participants placed their right hand on the keyboard (Fig. 1a). Their task was to counter the mechanical finger perturbation as quickly as possible. Task-related visual stimuli were presented on a computer monitor. At the start of each trial, participants were presented with a probability cue consisting of two vertical bars indicating the probability that either the index or ring finger would receive the perturbation (i.e., preparation phase). The probabilities were 0:100%, 25:75%, 50:50%, 75:25%, or 100:0% (index:ring) and were shown for the entire preparation phase. During this phase of the task, participants received continuous force feedback from two horizontal cursors projected on the screen. The cursors moved upward when the corresponding key was pressed. Participants were instructed to keep both cursors within a grey rectangular hold area, corresponding to a small pressure of 0.1-0.6N, throughout the preparation phase.

The trial then proceeded depending on whether it was a Go or No-Go trial. In Experiment 1, we used 2/3 Go and 1/3 No-Go trials. Experiment 2 included Go trials only. In Go trials, the duration of the preparation phase was randomly drawn from a uniform distribution between 1.5s and 2.5s. At the end of the preparation phase, the piston underneath one of the two fingers was activated, applying an upward force of ∼3.5N for 3s (i.e., execution phase). The padded clamp above the finger limited the upward movement of the finger to less than 5mm. Participants were instructed to respond as quickly as possible by pressing the piston down and hold it until the piston was deactivated. The force feedback was frozen at perturbation onset. At the end of the trial, the vertical position of the two cursors remained fixed for 2s showing the average force exerted by each finger during the execution phase (i.e., feedback phase). Participants were instructed that the cursor showing the force feedback of the perturbed finger should be in the target force area, corresponding to an average force response in the range between 3.5–8.5 N. The cursor of the non-stimulated finger should remain within the grey hold area. This delayed feedback helped participants adjust the amount of force applied in response to the perturbation. In the training session (see Procedures), participants learned to produce the correct force output within a few trials.

In No-Go trials, the perturbation was not applied, and the probability cue remained visible for 5.5s (i.e., the longest preparation period, 2.5s, plus execution duration in Go trials, 3s). The continuous force feedback provided by the two cursors was frozen at 2.5s as in Go trials. At the end of the trial, the probability cue disappeared and the delayed force feedback was provided as in Go trials, showing the average force exerted in the previous 3s. Because no perturbation had occurred, in the feedback phase both cursors were expected to be within the grey hold area.

In Experiment 1, in the feedback phase of each trial participants also received a score based on their performance. In Go trials, the score was based on the reaction time, defined as the interval between perturbation onset and the time when the force generated by any of the two fingers exceeded 3.5N. The score was assigned using a staircase system: +3 points for reaction time below the 25^th^ percentile of the previous run, +1 between the 25^th^ and 75^th^ percentile, and 0 above the 75^th^ percentile. The thresholds used in the first run were 0.25s and 0.50s for all participants. A negative score (−1) was assigned if any finger exerted >1N isometric force before the perturbation. In No-Go trials, the score was +1 if the participant kept both cursors within the grey hold area throughout the trial, and –1 otherwise. No scoring was used in Experiment 2.

### Procedures

In Experiment 1, participants completed a single fMRI session, consisting of 10 functional runs of 30 trials each. We used an event-related design. Each functional run included 6 trials for each probability cue (0:100%, 25:75%, 50:50%, 75:25%, 100:0%, index:ring). Of these, 4 were Go trials and 2 No-Go trials. In Go trials, the stimulated finger was randomly drawn from the cued probability distribution, such that the 100:0% and 0:100% cues were always followed by index and ring stimulation, respectively (see Supplementary Materials 1). Three 12.5-s periods of rest were randomly included in each functional run to allow for the estimation of baseline activation.

The day before the fMRI experiment, participants completed a short training session of 3-5 functional-like runs to familiarise themselves with the task. The training session was carried out in a mock fMRI scanner so that participants could familiarise themselves with the posture in which they would perform the task in the fMRI session.

Experiment 2 included Go trials only, resulting in 8 trial types. Because the task was performed while sitting at a desk, it was not necessary for participants to get accustomed to an unfamiliar posture. The training session was therefore replaced by a brief familiarisation with the task and equipment before starting the experiment.

### EMG recordings

In Experiment 2, we used an 11-channel surface EMG montage (Delsys, Trigno Research+ System, Trigno Duo Sensors) to record the activity of the extensor digitorum communis, extensor digiti minimi, extensor indicis, extensors of the thumb (extensor pollicis brevis and longus), flexor digitorum superficialis, abductor pollicis brevis, abductor digiti minimi, and first dorsal interosseous muscles. Raw EMG signals were acquired at 2148Hz. The skin was cleaned with alcohol before placing the electrodes for reducing impedance and improving the signal. We defined the ideal location of each electrode by asking the participant to perform slight isometric presses with each finger either in flexion or extension direction. The electrode was placed where muscle activation was maximal, as indexed by palpation and through continuous monitoring of EMG activity.

### Behavioural analysis

Visual inspection of the forces time-aligned to the perturbation suggested that the response scaled with the cued probabilistic predictions (see Fig. 1b). To quantify this observation, we averaged the isometric force produced by the stimulated finger (index or ring) in Go trials between 0.2-0.4s after perturbation onset (see dashed rectangles in Fig. 1b) and tested the within-participant effect of the cued probability using a repeated-measures ANOVA. This analysis was performed separately for index and ring finger perturbation.

We also hypothesised that, when the perturbation was applied to the finger cued with lower probability following the 25:75% or 75:25% cues, participants may initially respond with the finger cued with higher probability (75%) and then correct their response as sensory evidence accumulated. To quantify these corrections, we considered the 2-dimensional force trajectory (*f*_!_) generated by the index and ring finger in each trial between 0-0.5s after perturbation onset and defined the ideal straight response line (*c*) as the vector connecting the extreme points of this trajectory. We then computed, at each time point, the Euclidean distance of the force vector from this ideal straight line and averaged this distance over time (see Fig. 1d):

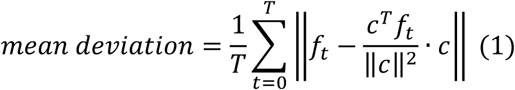

where *t* is the time relative to the perturbation onset in 2-ms time steps, and T=250 is the total number of time steps. If the force response unfolded along the ideal straight line, the mean deviation would be 0. On the other hand, if the response was initiated with the non-stimulated finger and later corrected to the stimulated finger, the force trajectory would deviate from the ideal straight line, thus increasing the mean deviation. For each stimulated finger, we then performed a paired-sample t-test between conditions in which the perturbation was delivered to the finger cued with lower probability, and all the other conditions.

### EMG preprocessing

The raw EMG signals were rectified, time-aligned to the perturbation and baseline-corrected by subtracting the mean EMG activity in the 0.1s preceding the perturbation. We defined four time windows relative to the perturbation, i.e., preparation (Pre, –0.1-0s), short-latency stretch reflex (SLR, 0.025-0.05s), long-latency stretch reflex (LLR, 0.05-0.1s) and voluntary response (Vol, 0.1-0.5s). Multivariate analysis and model fitting (see “Multivariate dissimilarity analysis” below) were performed separately on the mean EMG activity within each time window in each participant and condition.

### Imaging data acquisition

In Experiment 1, we used a 7T Siemens Magnetom scanner with a 32-channel head coil to acquire high-field fMRI data. The anatomical T1-weighted scan was acquired at the end of the scanning session using a Magnetization-Prepared Rapid Gradient Echo sequence (MPRAGE) with voxel size of 0.75 x 0.75 x 0.75mm isotropic (field of view = 208 x 157 x 110mm [A-P; R-L; F-H], encoding direction coronal). For functional scans (336 volumes) we used the following sequence parameters: GRAPPA 3, multiband acceleration factor 2, repetition time (TR) = 1.0s, echo time (TE) = 20ms, flip angle (FA) = 30°, slice number: 57, voxel size: 1.8 x 1.8 x 1.8mm isotropic. To estimate and correct for magnetic field inhomogeneities, we also acquired a gradient echo field map with the following parameters: transversal orientation, field of view: 210 x 210 x 160mm, 64 slices, 2.5mm thickness, TR = 475ms, TE = 4.08ms, FA = 35°.

### Preprocessing of fMRI data and general linear model

The preprocessing of functional images was performed with SPM12 (https://www.fil.ion.ucl.ac.uk/spm/) and custom MATLAB code and involved the following steps: (1) correction of geometric distortions using field maps^45^; (2) rigid-body motion realignment of all images to the first image of the first functional run; and (3) coregistration to the anatomical scan. No smoothing or normalisation to a standard template was applied at this stage.

We then analysed the pre-processed images using a general linear model (GLM)^46^, with separate regressors for preparation and execution. Preparation was modelled with five delta-function regressors, one for each probability cue, active at cue presentation in both Go and No-Go trials. Execution was modelled with eight delta-function regressors, active at perturbation onset in Go trials. This resulted in 13 regressors per run, plus an intercept. Each regressor was convolved with a two-gamma haemodynamic response function (HRF). We used a gridsearch approach to adjust the time to peak (4, 5, 6, 7, 8, and 9s) and the time to undershoot (10, 12, 14, 16, 18 and 20s) of the HRF and obtain the best fit to the BOLD timeseries. Before GLM estimation, the BOLD time series were high-pass filtered with a standard cutoff frequency of 128s. The 1^st^-level GLM analysis resulted in activation images consisting of the fitted beta coefficients across voxels for each of the 13 trial types, for each run and participant.

We also designed a separate GLM with preparation regressors only active in No-Go trials. This allowed us to obtain preparatory activity estimates that were not contaminated by execution activity. We then used these preparatory activity estimates to validate the multivariate analysis conducted on preparatory activity estimated using both Go and No-Go trials (see Supplementary Materials 3).

### Surface reconstruction and regions of interest definition

We used Freesurfer^47,48^ to reconstruct the white–grey matter and pial surfaces from each participant’s anatomical image. Each individual surface was then inflated to a sphere and nonlinearly aligned to the Freesurfer average atlas by matching cortical folding patterns, using sulcal depth and surface curvature to guide the alignment of gyri and sulci. Next, we resampled both hemispheres of each participant into a symmetric fs32k template, which represented each hemisphere using 32k vertices. In this way, by selecting corresponding vertices in each participant, it is possible to compare similar cortical regions.

We used a searchlight approach to assess the information represented across the entire cortical surface. A “searchlight” included all the voxels between the white-grey matter and pial surface within a ∼20-mm diameter circular region centred on a vertex of the fs32k template. We then fitted the relative weight of each representation of interest (see “Pattern component modelling”) to the beta coefficients estimated in the 1^st^-level GLM within each searchlight. Then, we assigned the resulting weights to the centre vertex.

We used a probabilistic cytoarchitectonic atlas projected onto the group surface^48^ to define seven anatomical ROI encompassing primary sensorimotor regions. M1 was defined by including all nodes belonging to Brodmann area 4 (BA4) within 2 cm from the hand knob^49^. S1 was defined by selecting the nodes belonging to BA1, 2 and 3 within 2 cm of the hand knob. We divided BA6 into a medial part (supplementary motor area, SMA), a lateral dorsal part (dorsal premotor cortex, PMd), and a ventral part (ventral premotor cortex, PMv). Finally, we separated the anterior and posterior parts of the superior parietal lobule (SPLa and SPLp) approximately at the midpoint of the intraparietal sulcus. To avoid contamination of activity between different ROI across sulci, we excluded voxels with more than 10% of their volume lying in a neighbouring ROI.

### Univariate analysis of fMRI data

To evaluate cortical activation during preparation and execution, we performed univariate contrasts of brain activity as compared to rest. The amount of activation was expressed as percent of signal change (PSC):

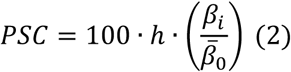

where ℎ is the median of the peak values of the regressor for condition *i* across runs, *β_i_* is the estimated regression coefficient for condition *i* and 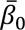 is the mean intercept across runs. We projected the PSC values calculated in each participant to the group surface via the individual reconstructed surfaces. For visualization purposes, we averaged the activation estimated across conditions, runs and participants, separately for preparation and execution (see Fig. 2c,d). For statistical testing, we conducted a one-sample t-test of the average activation values in each ROI against 0 across participants.

### Multivariate dissimilarity analysis

We used representational similarity analysis (RSA)^19–21^ to assess task-related representations in BOLD (i.e., beta coefficients from the 1^st^-level GLM) and EMG activity patterns. Within each ROI or searchlight, we first performed a multivariate spatial pre-whitening on the neural activity patterns using the residuals from the 1^st^-level GLM:

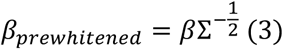

where *β* is the N (conditions x runs) by P (voxels) matrix of the estimated beta coefficients and Σ is the P-by-P noise covariance between voxels estimated from the residuals of the 1^st^-level GLM. The noise covariance Σ was regularised using the Ledoit-Wolf procedure^50^ to ensure invertibility. Because voxels often show different (and correlated) levels of noise, the weighting of activation patterns by the inverse noise covariance makes dissimilarity estimates more reliable^22^.

We used a similar pre-whitening procedure for EMG responses in the preparation, SLR, LLR and Vol time windows. For each time window and acquisition run, we first calculated the residuals of the EMG pattern observed in each condition relative to the mean across all conditions. We then used the variance of the residuals (*σ*^2(*w*)^) estimated separately for each channel to perform a univariate pre-whitening:

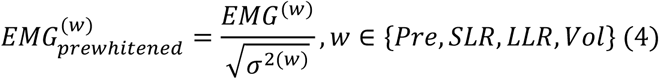

where *EMG*^(*w*)^ is the N-(conditions x runs)-by-P (channels) matrix of the EMG activity in the *w* time window. For both fMRI and EMG data, we then calculated the cross-validated squared Mahalanobis (crossnobis) dissimilarity *d* between conditions *i* and *j* as follows:

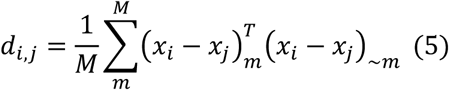

where *M* is the number of runs, *x_i_* and *x_j_* are the pre-whitened (neural or EMG) activity patterns for conditions *i* and *j*, either from run *m* or averaged across all the other runs (∼*m*). Cross-validation makes the dissimilarity estimates unbiased by measurement noise^22^. Because measurement noise pulls activity patterns in random directions, without cross-validation the expected dissimilarity between two activity patterns would be always larger than 0, even when they are identical and only differ by their noise. With cross-validation, the expected dissimilarity between identical patterns is 0, which means that we could test the average dissimilarity against 0 using a one-sided t-test to determine whether activity patterns carried reliable information. Note that, especially when two patterns are very similar, cross-validated dissimilarity estimates can be negative^19,20,51^.

### Contribution of finger pre-activation to the expectation representation

We assessed finger pre-activation by calculating the difference between index and ring finger force during the preparation phase (see Fig. 3g). We then performed two control analyses to ensure that the expectation representation in M1 and S1 during preparation was not driven by subtle finger pre-activation.

First, we calculated the average crossnobis dissimilarity between the mean 5-finger force patterns for each probability cue measured in the preparation phase. Then, we performed a linear regression analysis between the average force dissimilarity (see Fig. 3h) and the average BOLD (see Fig. 3c) dissimilarity observed during preparation in each participant. Two separate regressions were conducted using the average BOLD dissimilarity from M1 and S1. If the dissimilarities between BOLD activity patterns were solely driven by finger preactivation, then the intercept of the regression should not be significantly larger than 0; that is, participants that did not show finger pre-activation should also not show any BOLD dissimilarity. To test this, we performed a one-sided t-test of the intercept estimate against 0.

In the second analysis, we regressed out the pairwise force dissimilarities between conditions (see Fig. 3h) from the pairwise BOLD dissimilarities in each ROI (see Fig. 3c) and within each participant. Then, we tested the residual neural dissimilarities against 0 to evaluate the residual encoding and correlated them with the RDMs for the expectation and uncertainty models (see Fig. 3a) to assess the residual information content (see Supplementary Materials 4).

### Pattern component modelling

We used pattern component modelling (PCM)^52^, a complementary framework to RSA, to characterise the nature of the representational geometry in neural and muscle activity patterns. Rather than estimating and evaluating dissimilarities between activity patterns, PCM is a probabilistic framework that evaluates the marginal likelihood of the observed activity patterns under the hypothesis that they follow a multivariate Gaussian distribution with mean 0 and covariance matrix *G*. Because we assumed a mean of 0, we did not subtract out the mean of each condition across voxels, thus *G* is more accurately defined as the second moment matrix of the distribution. The second moment matrix can be directly translated to squared Euclidean distances (*D*) through the equation:

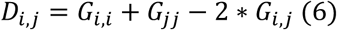

where *i* and *j* are two different conditions (e.g., two different probability cues or stimulated fingers in the current experimental design). This allows us to visualise the representational models as RDMs (see Fig. 3a and 6a), while still using the more powerful^19^ approach of PCM for model evaluation. Note that the squared Euclidean distance is mathematically equivalent to crossnobis dissimilarity when the activity patterns are pre-whitened^22^.

### Representational models for preparation

For the preparation phase we considered two different representations of the probability cue. The expectation representation predicted that the activity of each voxel scaled linearly with a feature vector (*f*) corresponding to the difference in probability between index and ring:

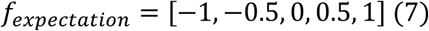

The uncertainty representation predicted that the activity patterns scaled with the variance of a Bernoulli distribution, defined as:

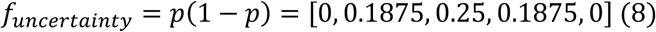

where *p* is the probability of a certain outcome (e.g., index stimulation) and 1 − *p* is the probability of the opposite outcome (e.g., ring stimulation).

For each representational model, the predicted second moment matrix *G* was defined as the outer product of *f*:

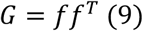

For visualisation, we then calculated the predicted dissimilarity matrices of each model from the corresponding second moment matrices according to Eq. 6 (see Fig. 3a and 6a).

### Representational models for execution

For execution, we designed three different representational models that predicted the neural or EMG response in 8 conditions, including 4 different probability levels (i.e., 25% to 100%) for each finger. The sensory input representation predicted that activity patterns were simply modulated based on the identity of the stimulated finger and was defined as:

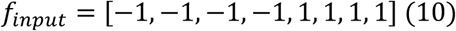

with −1 and 1 indicating index and ring finger, respectively.

We also assessed the expectation representation, which predicted that activity patterns scaled with the difference between index and ring finger probability:

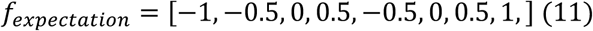

Finally, the surprise representation predicted that the activity patterns scaled linearly with the Shannon surprise, defined as the negative log-probability of the observed event given the cue:

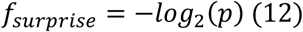

where *p* = [1, 0.75, 0.5, 0.25, 0.25, 0.5, 0.75, 1], i.e., the probability cued on the stimulated finger. Note that the Shannon surprise reflects the absolute prediction error between finger and cue.

### Model fitting and evaluation

In PCM, the predicted second moment matrix (*Ĝ*) of the true activity patterns underlying the observed data is modelled as the linear combination of different representational models (*G*_h_):

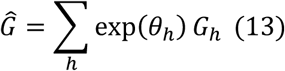

where exp (*θ_h_*) is the weight parameter of the ℎ*^th^* representational model, and *G_h_* is the predicted second moment matrix. Because weights cannot be negative, using the exponential allows for unconstrained optimisation of the model parameters. To make the weights comparable, we normalised the trace of the predicted second moment matrix *G_h_* for each representational model to 1. In this way, the weight can be used to estimate the amount of variance explained by each representation included in the model. To model the overall second moment matrix of the observed data across all runs, PCM also estimates the variance of the measurement noise. Importantly, this approach relies on the assumption that each representation spans an independent neural dimension. Therefore, the weight of each representation reflects how strongly the corresponding information is encoded in a certain brain region independently from all other representations in the model.

For the fMRI data, we fitted the relative weights of each representational model to the pre-whitened beta coefficients, separately for each task phase (preparation and execution) and within each ROI or searchlight. We performed one-sided dependent-sample t-tests to compare the relative weight of different representations within the same ROI. Then, to directly compare the information encoded in different premotor-parietal vs. M1-S1, we averaged the weight of each representation within each ROI group and performed a one-sided de-pendent-sample t-test on the weight of a representation of interest (e.g., expectation in Fig. 3e and sensory input in Fig. 6d) relative to the sum of all representations of interest (e.g., expectation+uncertainty in Fig. 3e and sensory input+surprise in Fig. 6d).

To assess the expectation (and uncertainty) representation in the LFPs and spiking activity recorded from non-human primates in PMd and S1, we time-aligned the recordings to cue presentation and perturbation onset and then fitted the relative weight of each representation at each time point (and frequency band, for the LFPs). Then, to assess whether the expectation encoding differed across areas (i.e., PMd vs. S1) and recording modality (i.e., spiking activity vs. LFPs), we averaged the expectation weight over 0.64s (and between 10-20Hz for LFPs) after cue presentation (see grey-shaded time interval in Fig. 4b) and performed a 2-by-2 ANOVA. Because spiking activity and LFPs are recorded in different units, it is not possible to directly compare the expectation weight in the two recording modalities. For this reason, we normalised the expectation weight of each recording modality over the variance of the measurement noise before performing the ANOVA. Fig. 4a,b and Fig. S6 display the expectation and uncertainty weights normalised by the measurement variance.

To establish whether a predicted representation contributed significantly to explain activity, we first fitted all the possible combinations of the candidate representations (e.g., expectation alone, uncertainty alone and expectation+uncertainty). Then, we calculated the log-Bayes factor (*BF_F_*) of each representation, defined as the difference between the marginal log-likelihoods of the activity patterns under the models that included the representation and those that did not:

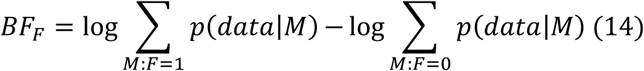

Where *F* is the representation of interest (i.e., expectation, uncertainty, sensory input, or surprise) and *M* is a model including a combination of representations (*F* = 1 indicates that the representation of interest is included in *M*). The maximum likelihood of the observed activity under model *M* is given by *p*(*data*|*M*) and was estimated using the Akaike Information Criterion (AIC), which corrects the maximum likelihoods for the model complexity^53^.

For fMRI data, we tested the log-Bayes factor of each representational model against 0 using a one-sample t-test across participants in each ROI. A positive log-Bayes factor indicates that the representational component helped to explain the activity patterns in the context of all the other components.

For the LFPs and spiking activity, we used cluster-based permutations to establish the significance of the log-Bayes factor in different time (and frequency) bins, while controlling for multiple comparisons^54^. First, we calculated a one-sample t-statistic against 0 across sessions and defined clusters of contiguous significant time(-frequency) bins. We then generated a null distribution by randomly inverting the sign of the log-Bayes factor for a random subset of sessions and recomputing the t-statistic over 1,000 permutations. In each permutation, we thresholded the t-values at the uncorrected p<0.05 level and recorded the cluster of contiguous significant bins with the largest weighed size, defined as the number of bins in the cluster multiplied by the sum of the absolute t-values within the cluster. The observed clusters were considered significant if their weighted size exceeded the 95th percentile of the null distribution.

### Correlation model

In Experiment 1, we used a PCM correlation model to obtain maximum-likelihood correlation estimates between BOLD activity patterns corrected for the attenuation bias induced by measurement noise. First, to assess how expectation patterns were related to those elicited by the sensory input, we calculated, in each run, the difference between BOLD activity patterns for index (i.e., 100:0% and 75:25% cues) vs. ring (0:100% and 25:75% cues) expectation in the preparation phase (i.e., expectation pattern). Similarly, we calculated the difference between BOLD activity patterns for index vs. ring perturbation in the execution phase (i.e., sensory input pattern). Then, we estimated the maximum-likelihood correlation between the expectation and sensory input patterns. The correlation model uses the repeated measures of the two patterns (*x* and *y*) across runs to estimate their signal variances (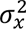 and 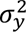) as well as the variance of independent Gaussian noise 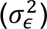. In this way, the maximum-likelihood correlation estimate (*ρ*) reflects the true correlation of the activity patterns underlying their noisy run-wise estimates. Specifically, the predicted second moment matrix (*Ĝ*) of the true activity patterns is modelled as:

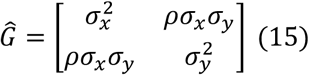

We obtained the maximum-likelihood correlation estimates between the true activity patterns both for each individual participant and for the group (see black dots and dashed horizontal red line, respectively, in Fig. 5). To obtain confidence intervals for the group estimate, we conducted a bootstrap procedure, resampling the participants with replacement at each iteration. We performed 1000 iterations and calculated the 95% central confidence interval of the correlation estimate (see dashed grey areas in Fig. 5).

We used a similar approach to estimate the correlation between the pattern components for expectation and sensory input in BOLD, EMG, LFPs and spiking activity during execution. In this case, we calculated the difference between execution activity following the 75-25% vs. 25-75% probability cues (i.e., expectation pattern) and between index vs. ring finger (or flexion vs. extension, for LFPs and spiking activity in monkeys) perturbation (i.e., sensory input pattern). Then, we obtained maximum-likelihood estimates of the correlation between the expectation and sensory input patterns using PCM. For the EMG data, we estimated the correlation using the pre-whitened (see Eq. 3) mean activity patterns in the preparation, SLR, LLR and Vol time windows. For the same correlation in LFPs and spiking activity, we used the pre-whitened (as done for EMG, see Eq. 3) mean patterns across electrodes (LFPs) or units (spiking activity) between 40-240ms after the perturbation. For the LFPs, separate correlations were estimated in the alpha (8-13Hz), beta (13-25Hz) and gamma (25-100Hz) frequency bands.

To diagnose the reliability of our correlation estimates, we plotted the correlation estimates against the estimated functional signal-to-noise ratio (fSNR) of the two contrasts, defined as:

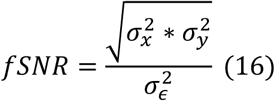

At low fSNR, maximum-likelihood correlation estimates become unstable and may fall at the parameter bounds (i.e., correlation=±1). For example, in the preparation phase of EMG recordings (see Fig. 7f), the fSNR was close to 0 and the preparation-execution correlation estimate was −1 for the majority of the participants. This scenario flags the correlation estimate as unreliable.

### Low-dimensional projections of BOLD activity patterns

We used a multidimensional scaling approach to obtain the low-dimensional projections of BOLD activity shown in Fig. 8a. This corresponds to performing an eigen-decomposition of the second moment matrix of the observed activity patterns, yielding orthogonal dimensions in voxel space that capture most of the variance across conditions. In this way, the projection of the activity patterns onto the first two principal components preserves the dominant representational geometry of the second moment matrix and can be used for visualisation.

### Electrophysiological recordings in non-human primates

The spiking data from the non-human primate electrophysiological datasets are publicly available^23^ and described in detail in a recently published paper^9^. In brief, monkeys countered mechanical perturbations delivered with a KINARM robot exoskeleton (BKIN Technologies)^55^ that could rotate the elbow either in flexion or extension direction. Before the perturbation, the monkeys received probabilistic information about the upcoming perturbation (0:100%, 25:75%, 50:50%, 75:25%, 100:0%, flexion:extension). Electrophysiological recordings were carried out using high-density Neuropixels probes (1.0 - 1 cm, 1.0 NHP - 1 cm, and 1.0 NHP - 4.5 cm), pre-processed using a custom pipeline (https://github.com/JonathanAMichaels/PixelProcessingPipeline), and spike sorted with Kilosort 2.0^56^. The LFPs were read from the Neuropixels LF stream, which was recorded at 2,500Hz. We downsampled the initial 384 channels to a total of 32 channels and then performed a time-frequency analysis using the FieldTrip toolbox^57^. We defined fifty frequencies of interest logarithmically (log10) spaced from 1 Hz to 400 Hz. The time bins of interest were sampled at a resolution of 0.01 seconds. After the LFPs were demeaned and bandpass filtered (1-400 Hz, 3rd order), we calculated the power spectrum in each time bin using the multi-taper convolution method with a Hanning taper. The time windows for the convolution were dynamically adjusted relative to the frequencies of interest to cover 5 cycles at each frequency. For the subsequent analysis, we pooled the recording sessions from both monkeys (PMd, 17 sessions; M1, 9 sessions; S1, 9 sessions) totalling ∼35000 trials overall.

## Acknowledgements

Experiments were funded by a project grant from the Canadian Institutes of Health Research (CIHR, PJT – 175010) to JD and AP, a Discovery grant from the Natural Sciences and Engineering Research Council of Canada (NSERC, RGPIN-2016-04890) to JD, and the Canada First Research Excellence Fund (BrainsCAN) to Western University. AP received a salary award from the Canada Research Chairs Program.

