## Supplementary Materials for "Sensory expectations and prediction error during feedback control in the human brain"

### Supplementary Materials 1: Experimental design

Table S1. Number of trials for each condition (i.e., probability cue and perturbed finger) in each functional run in Experiment 1. Experiment 2 had the same design but without No-Go trials.

| **Condition** | | **Perturbed finger** | | |
|---|---|---|---|---|
| | | Index | Ring | No-Go |
| **Probability cue** | 100-0% | 4 | 0 | 2 |
| | 75-25% | 3 | 1 | 2 |
| | 50-50% | 2 | 2 | 2 |
| | 25-75% | 1 | 3 | 2 |
| | 0-100% | 0 | 4 | 2 |

### Supplementary Materials 2: Right hemispheric results

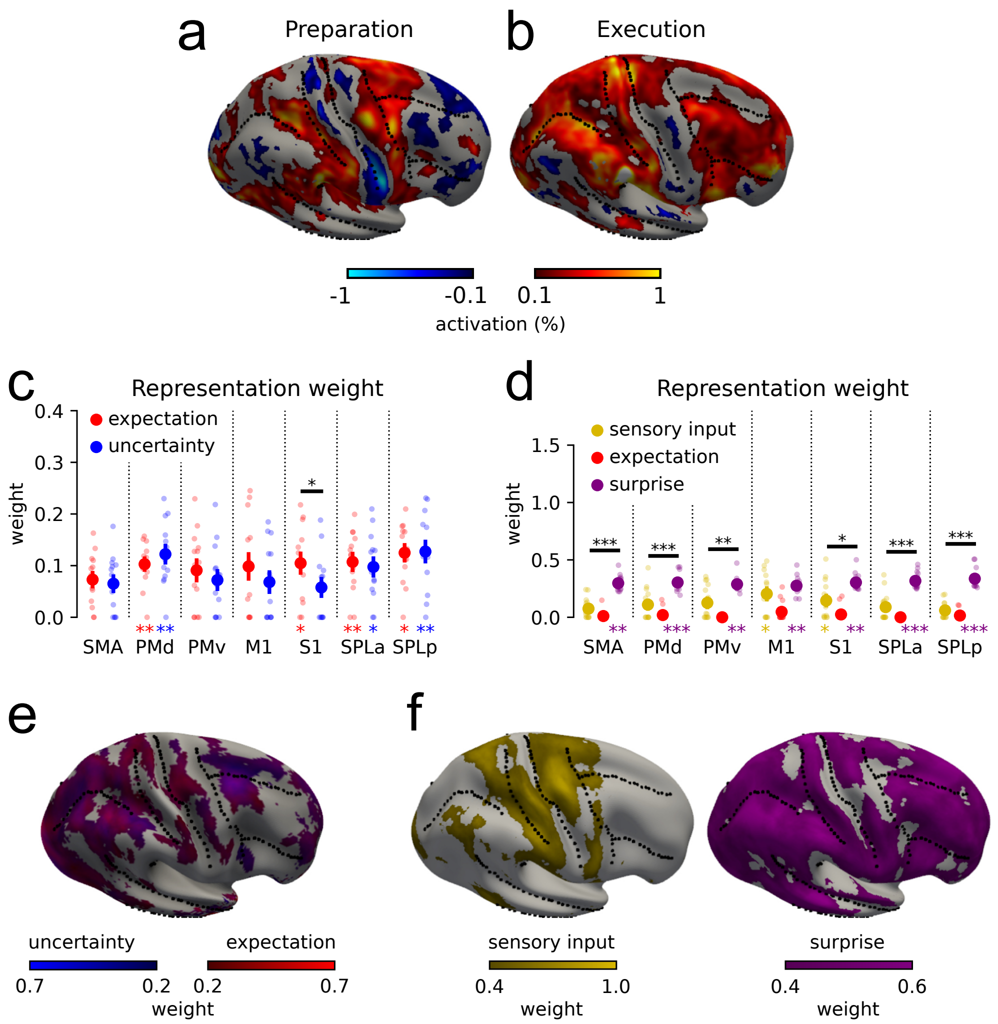


Figure S2. Activation relative to resting baseline during **(a)** preparation and **(b)** execution in the right hemisphere. Relative weight of **(c)** expectation and uncertainty during preparation and **(d)** sensory input, expectation and surprise during execution. Dots denote individual participant data. Asterisks below the bars indicate that the representation had a log-Bayes factor significantly larger than 0 across participants. The horizontal bars denote significant weight differences between expectation and uncertainty (panel c) and between sensory input and surprise (panel d). Error bars denote S.E.M. across participants. **(e)** Weight of expectation and uncertainty component during preparation in a continuous searchlight analysis conducted on the surface of the right hemisphere. **(f)** Searchlight analysis for sensory input and surprise during execution across the right hemisphere.

Table S2. ROI-based statistics for the right hemisphere. See Table 1 in the main text for statistical tests and conventions.

| Phase | Statistics | | SMA | PMd | PMv | M1 | S1 | SPLa | SPLp |
|---|---|---|---|---|---|---|---|---|---|
| Preparation | Activity>rest | | $t_{13}$=2.922, P=0.006 | $t_{13}$=2.965, P=0.006 | $t_{13}$=2.804, P=0.008 | $t_{13}$=0.846, P=0.206 | $t_{13}$=-0.380, P=0.645 | $t_{13}$=1.956, P=0.036 | $t_{13}$=0.921, P=0.187 |
| | Encoding | | $t_{13}$=3.396, P=0.002 | $t_{13}$=5.916, P<0.001 | $t_{13}$=3.119, P=0.004 | $t_{13}$=4.753, P<0.001 | $t_{13}$=2.659, P=0.010 | $t_{13}$=5.338, P<0.001 | $t_{13}$=5.531, P<0.001 |
| | log-Bayes factor | Expectation | $t_{13}$=0.429, P=0.337 | $t_{13}$=2.837, P=0.007 | $t_{13}$=1.484, P=0.081 | $t_{13}$=1.660, P=0.060 | $t_{13}$=2.043, P=0.031 | $t_{13}$=3.009, P=0.005 | $t_{13}$=2.600, P=0.011 |
| | | Uncertainty | $t_{13}$=0.426, P=0.338 | $t_{13}$=2.967, P=0.005 | $t_{13}$=0.871, P=0.200 | $t_{13}$=0.962, P=0.177 | $t_{13}$=0.845, P=0.207 | $t_{13}$=1.789, P=0.048 | $t_{13}$=2.861, P=0.007 |
| Execution | Activity>rest | | $t_{13}$=2.816, P=0.007 | $t_{13}$=2.061, P=0.030 | $t_{13}$=1.298, P=0.109 | $t_{13}$=1.049, P=0.157 | $t_{13}$=3.436, P=0.002 | $t_{13}$=2.994, P=0.005 | $t_{13}$=5.307, P<0.001 |
| | Encoding | | $t_{13}$=6.603, P<0.001 | $t_{13}$=4.902, P<0.001 | $t_{13}$=3.899, P=0.001 | $t_{13}$=4.887, P<0.001 | $t_{13}$=4.371, P<0.001 | $t_{13}$=3.844, P=0.001 | $t_{13}$=9.826, P<0.001 |
| | log-Bayes factor | Sensory input | $t_{13}$=-0.699, P=0.752 | $t_{13}$=1.483, P=0.081 | $t_{13}$=1.150, P=0.135 | $t_{13}$=1.910, P=0.039 | $t_{13}$=1.589, P=0.068 | $t_{13}$=0.537, P=0.300 | $t_{13}$=0.033, P=0.486 |
| | | Expectation | $t_{13}$=-17.804, P=1.000 | $t_{13}$=-14.386, P=1.000 | $t_{13} \ll 0$, P=1.000 | $t_{13}$=-2.021, P=0.968 | $t_{13}$=-7.424, P=1.000 | $t_{13} \ll 0$, P=1.000 | $t_{13}$=-16.485, P=1.000 |
| | | Surprise | $t_{13}$=3.274, P=0.003 | $t_{13}$=4.673, P<0.001 | $t_{13}$=3.530, P=0.002 | $t_{13}$=3.375, P=0.002 | $t_{13}$=3.231, P=0.003 | $t_{13}$=3.959, P<0.001 | $t_{13}$=4.047, P<0.001 |

### Supplementary Materials 3: Preparatory activity in No-Go trials

Because the BOLD response is slow and the perturbation was applied only 1.5-2.5s after cue presentation, there was a risk that preparatory activity estimates were contaminated by execution-related activity. To rule out that the representational geometry during preparation is the result of such contamination, we used a separate GLM where the 5 preparation regressors were estimated using No-Go trials only. Each regressor was a boxcar function active for 2s following cue presentation. This GLM also included a separate 2-s boxcar regressor active during cue presentation in Go trials and a delta-function regressor that was active at perturbation onset (totalling 7 regressors overall). All regressors were convolved with a haemodynamic response function (HRF), with response delay and undershoot delay optimised in each participant to obtain the best fit with the BOLD timeseries.

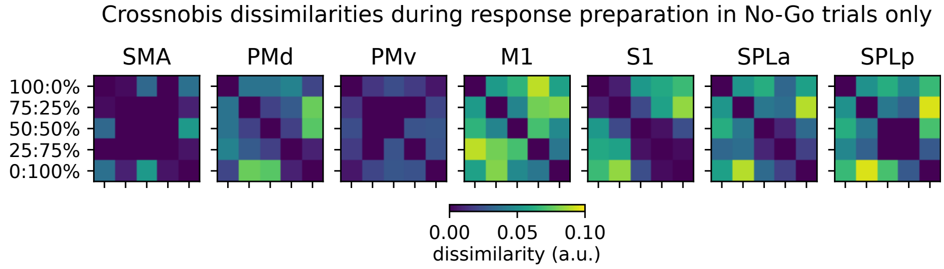


Figure S3. Crossnobis dissimilarities estimated from preparatory activity in No-Go trials only.

In each ROI, we calculated the crossnobis dissimilarities between the estimated preparatory activity patterns from No-Go trials only (Fig. S3) and correlated them with those estimated using the main GLM design in each participant (see Fig. 3c). All correlations were significantly larger than 0 across participants (all $\rho>0.378$, all $t_{13}>4.088$, all $Ps<0.001$, one-sided t-test).

We concluded that the information content in the preparation phase remains similar when preparatory activity is estimated from both Go and No-Go trials and from No-Go trials only.

### Supplementary Materials 4: Residual BOLD dissimilarities after regressing out pairwise force dissimilarities during preparation

To provide further support to the conclusion that the expectation representation during the preparation phase in M1 and S1 is not driven by subtle finger pre-activation, we regressed the pairwise BOLD dissimilarities between conditions in each ROI (see Fig. 3c) onto the corresponding pairwise force dissimilarities observed in this phase (see Fig. 3h). We then tested the residual neural dissimilarities against 0 to evaluate whether the encoding of the probability cue survived (one-sided t-test). We found that the residual BOLD dissimilarities (Fig. S4a) remained significantly positive across all ROI (Table S4, first row).

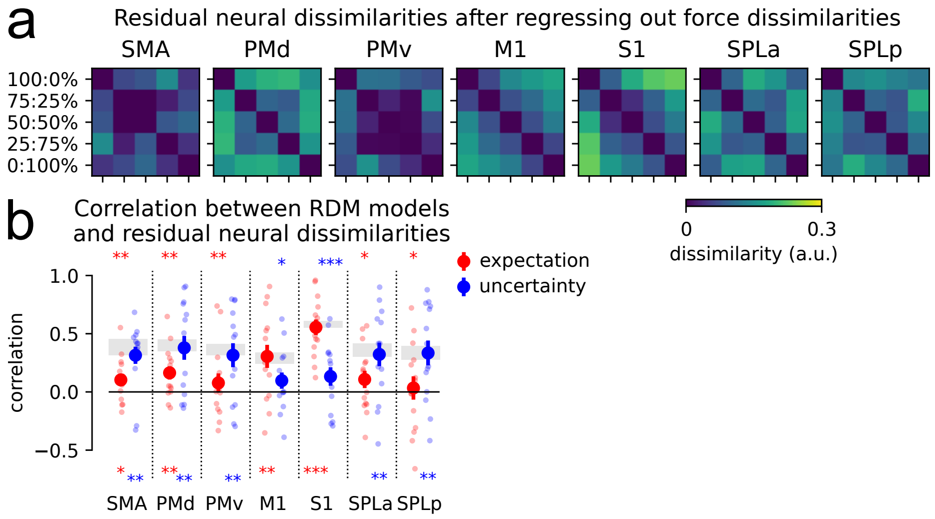


Figure S4. **(a)** Residual BOLD dissimilarities after regressing out pre-activation force dissimilarities. **(b)** Correlation between residual BOLD dissimilarities and the RDMs of the expectation and uncertainty models (see Fig. 3a in the manuscript). The shaded area denotes the lower and upper bounds of the noise ceiling. Bottom asterisks show significance against 0. Top asterisks show significance against the lower bound of the noise ceiling (*P<0.05, **P<0.01, ***P<0.001). Error bars denote ±SEM across subjects.

We then correlated the RDMs for the expectation and uncertainty models (see Fig. 3a) with the residual neural dissimilarities to assess the residual information content. The average correlation across participants in each ROI was tested against 0 and against the lower bound of the noise ceiling (one-sided t-test). The noise ceiling was defined as the correlation between the dissimilarities in each participant and the group mean. For the upper bound, the group mean included all participants; for the lower bound, the participant being evaluated was removed from the group mean (i.e., leave-one-out cross-validated reliability). The expectation and uncertainty models explained the residual BOLD dissimilarities up to noise ceiling in M1-S1, and in premotor-parietal areas, respectively. Therefore, the information content remained the same after the force dissimilarities were regressed out from the BOLD dissimilarities, consistent with a genuine (i.e., not caused by finger pre-activation) expectation representation during preparation.

Table S4. ROI-based statistics for preparation after regressing out the force dissimilarities from neural dissimilarities at individual-participant level. T-values are one-sided t-tests against 0 or lower noise ceiling with uncorrected P-values provided.

| Statistics | | SMA | PMd | PMv | M1 | S1 | SPLa | SPLp |
|---|---|---|---|---|---|---|---|---|
| **Encoding** | | $t_{13}$=2.887, P=0.006 | $t_{13}$=4.682, P<0.001 | $t_{13}$=3.494, P=0.002 | $t_{13}$=3.975, P=0.001 | $t_{13}$=4.531, P<0.001 | $t_{13}$=3.977, P=0.001 | $t_{13}$=5.146, P<0.001 |
| **Model correlations** | Expectation vs. 0 | $t_{13}$=1.960, P=0.036 | $t_{13}$=2.745, P=0.008 | $t_{13}$=0.908, P=0.190 | $t_{13}$=2.970, P=0.005 | $t_{13}$=8.051, P<0.001 | $t_{13}$=1.392, P=0.094 | $t_{13}$=0.324, P=0.376 |
| | Expectation vs. noise ceiling | $t_{13}$=-4.032, P=0.001 | $t_{13}$=-3.087, P=0.004 | $t_{13}$=-2.809, P=0.007 | $t_{13}$=0.597, P=0.720 | $t_{13}$=0.011, P=0.504 | $t_{13}$=-2.541, P=0.012 | $t_{13}$=-2.327, P=0.018 |
| | Uncertainty vs. 0 | $t_{13}$=4.113, P=0.001 | $t_{13}$=3.560, P=0.002 | $t_{13}$=2.952, P=0.006 | $t_{13}$=1.341, P=0.101 | $t_{13}$=1.607, P=0.066 | $t_{13}$=3.071, P=0.004 | $t_{13}$=3.010, P=0.005 |
| | Uncertainty vs. noise ceiling | $t_{13}$=-0.018, P=0.493 | $t_{13}$=0.292, P=0.613 | $t_{13}$=-0.001, P=0.500 | $t_{13}$=-2.029, P=0.032 | $t_{13}$=-5.109, P<0.001 | $t_{13}$=0.179, P=0.570 | $t_{13}$=0.532, P=0.698 |

### Supplementary Materials 5: Beta desynchronisation

In line with previous work[1–5], in the LFPs recorded from non-human primates the frequency band roughly spanning between ~8-30Hz (i.e., alpha- and beta-band) exhibited a marked desynchronisation during response preparation and execution (Fig. S5a). During execution, we also observed a sharp increase in the power of oscillations >30Hz (i.e., gamma power), again consistent with well-established modulations of cortical rhythms accompanying movement[6].

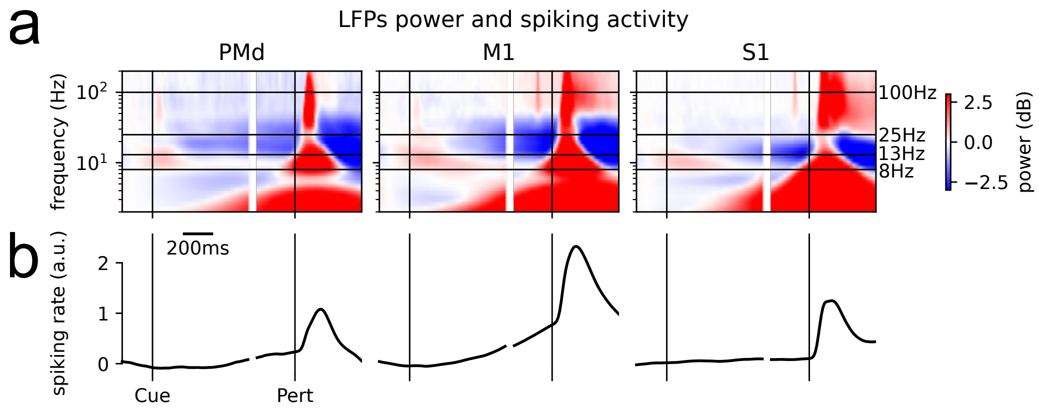


Figure S5. **(a)** LFP spectrogram and **(b)** spiking activity time-aligned to the probability cue (Cue) and perturbation onset (Pert) and averaged across sessions, conditions and channels/units.

### Supplementary Materials 6: Uncertainty representation in non-human primates

In non-human primates, the uncertainty representation was much weaker than expectation across all the examined regions in both LFPs (Fig. S6a) and spiking activity (Fig. S6b). However, the log-Bayes factor was significantly larger than 0 (P<0.05, cluster-based permutations) after cue presentation both in the spiking activity and LFPs recorded in PMd, suggesting that, like humans, also monkeys represented this information during preparation.

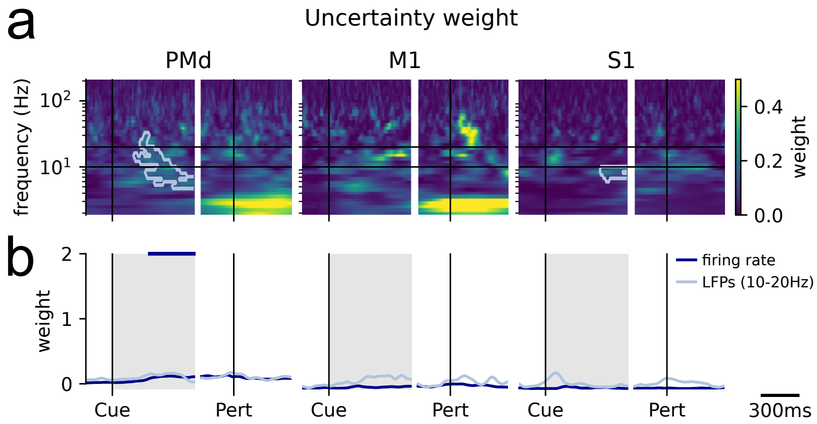


Figure S6. Weight of the uncertainty representations in **(a)** the LFPs and **(b)** spiking activity from PMd, M1 and S1 time-aligned to cue presentation (Cue) and perturbation onset (Pert). The light blue contours in panel a and the horizontal dark blue bars in panel b mark clusters where the log-Bayes factor was significantly larger than 0 in the LFPs and spiking activity, respectively. The scale used in this figure is the same used in Figure 4 for the expectation weight, to facilitate the comparison between the two representations.
